# Dopamine neurons promotes wakefulness via the DopR receptor in the Drosophila mushroom body

**DOI:** 10.1101/2020.04.29.069229

**Authors:** Margaret Driscoll, Steven Buchert, Victoria Coleman, Morgan McLaughlin, Amanda Nguyen, Divya Sitaraman

## Abstract

Neural circuits involved in regulation of sleep play a critical role in sleep-wake transition and ability of an organism to engage in other behaviors critical for survival. The fruit fly, Drosophila melanogaster is a powerful system for the study of sleep and circuit mechanisms underlying sleep and co-regulation of sleep with other behaviors. In *Drosophila*, two neuropils in the central brain, mushroom body (MB) and central complex (CX) have been shown to influence sleep homeostasis and receive neuromodulator input critical to sleep-wake switch.

Dopamine neurons (DANs) are the primary neuromodulator inputs to the MB but the mechanisms by which they regulate sleep- and wake-promoting neurons within MB are unknown. Here we investigate the role of subsets of DANs that signal wakefulness and project to wake-promoting compartments of the MB. We find that inhibition of specific subsets of PAM and PPL1 DANs projecting to the MB increase sleep in the presence of strong wake-inducing stimuli that reduces GABA transmission, although activity of these neurons is not directly modulated by GABA signaling. Of these subsets we find that DANs innervating the γ5 and β’2 MB compartments require both DopR1 and DopR2 receptors located in downstream Kenyon cells and mushroom body output neurons (MBONs). Further, we report that unlike the activity of wake-promoting MBONs and KCs, whose activity is modulated by sleep-need and PAM-DAN activity is independent of sleep-need. We have characterized a dopamine mediated sleep-circuit providing an inroad into understanding how common circuits within MB regulate sleep, wakefulness and behavioral arousal.

## Introduction

Sleep, wakefulness and arousal represent internal states that control multiple physiological and behavioral processes. Transition between these states and persistence of these individual states involves neural circuits that are dispersed throughout the brain and interact with systems involved in controlling hunger, goal-directed behavior and memory formation.

Sleep and wakefulness are regulated by two processes, known as process S and process C, where Process S is a homeostatic process by which sleep pressure increases during wakefulness and decreases during sleep, and process C is a circadian process that is controlled by circadian pacemaker circuits of the brain and determines the propensity to sleep during specific times of the day and night.^1-3^ The process S and C model has provided a critical framework but regulation of sleep, wakefulness and arousal goes beyond these processes. While, sleep provides many benefits, it limits the ability of the organism to engage in other behaviors critical for survival. Sleep and wakefulness are actively balanced and influenced by motivational or cognitive processes that modulate transition and persistence of these states^4,5^. Hence, there are additional processes that regulate sleep and are strongly influenced by conflicting needs (internal and external) which are not accurately represented in the existing models.

At the behavioral level motor output, attention, motivation, reward, and feeding all function on the basis of wakefulness and represent different levels of arousal and involve conserved neuromodulators like dopamine^6-8^, serotonin^9-12^ and histamine^13-16^. Neuromodulators are highly conserved across species and can target synapses by altering excitability of neurons that generate variable output from defined circuits^17-19^. Hence, a comprehensive understanding of how neuromodulators influence structures involved in sleep, wakefulness and other associated behaviors can provide an inroad into understanding the highly plastic nature of sleep regulation.

Dopamine is a wake-inducing neuromodulator in flies and mammals^20-24^. The ability to manipulate genes and genetically defined neural circuits has allowed identification of distinct sleep-wake microcircuits and their modulation by dopamine in flies and rodents^4,5,20-22,25-28^. Broad manipulations of DANs using genetic and pharmacological approaches shows that dopamine is required for wakefulness and this is further supported by increased sleep in receptors and transporters mutants^25,28^.

In the most well-defined dopamine-mediated wake circuit in flies, a cluster of PPL1 neurons project to the central complex innervating the dorsal fan-shaped body region (dfb) that has been shown to encode sleep need and critical for sleep homeostasis. Specifically, the dorsal fan shaped body dependent sleep switch is inhibited by dopamine input via DopR2 signaling and altered potassium conductance^26,29,30^. Artificial PPL1 cluster activation or direct dopamine application electrically silence sleep-promoting dfb neurons by altering the receptivity of the dopamine arousal signal, but it’s unclear if the PPL1 neurons themselves signal sleep-need^26,29,30^. While, DA dependent mechanisms of sleep-regulation have been identified within the CX^29^ and circadian clock system^31^, the widely used TH-GAL4 excludes several PAM neurons which provide key DA inputs to MBs^32^.

Targeted thermogenetic activation of DANs projecting to regions outside CX that innervate MB (PAM and PPL1) are also wake-promoting suggesting that distinct dopamine clusters induce wakefulness via distinct neural structures^3,33^. However, it is unclear if dopamine projections outside the CX are also involved in sleep homeostasis and if they use the same or distinct sleep-wake switching mechanisms.

Two separate clusters of neurons PAM and PPL (~ 130 neurons) account for the majority of dopamine signaling in the fly brain and represent the most extensive neuromodulator input to the MB, a key center required for associative learning in insects^34,35^.

About 2000 Kenyon cells, send out parallel axonal fibers that form the MB lobes with dendrites organized within the calyx. 22 MBONs innervate the lobes and receive input from the KC’s forming distinct compartments and each of these compartments receives modulatory input from one or more of the 20 subsets of dopaminergic neurons (DANs)^3,36-40^. The core KC-MBON circuits are modulated by DANs, which signal olfactory cues, satiety, wakefulness, negative and positive valence and novelty/familiarity of stimuli^40-46^. The current model of DAN regulation posits a model where PPL1 signal punishment or negative valence and PAM DANs signal reward^36,44,47-50^. But there is growing evidence that DANs signal a wide-range of information to the MB about novelty^40^, satiety^41^, locomotion^51,52^ and sleep/activity states^3,53^. While, there is neuroanatomical, physiological and biochemical evidence that DANs adjust and tune synaptic weights between KCs and MBONs, the mechanisms related to receptors and downstream signaling are not uniform across compartments^47,54,55^.

In our previous work, we comprehensively identified the KCs, MBONs, and DANs that control sleep, by performing an unbiased thermogenetic activation screen using a new library of intersectional split-GAL4 lines^37,56^. We identified several classes of sleep-controlling MBONs with dendrites in distinct lobe compartments: cholinergic sleep-promoting MBON-γ2α’1, and glutamatergic wake-promoting MBON-γ5β’2a/β’2mp/β’2mp_bilateral (MBON 01, 03, and 04) and MBON-γ4 > γ1γ2 (MBON 5). The sleep effects were consistent in both males and females and we did not find any sexually dimorphic sleep phenotypes or neuroanatomical differences in GFP expression^33,56^. We also determined that α’/β’ and γm KCs are wake-promoting and γd KCs are sleep-promoting, and that α’/β’ and γm KCs promote wake by activating MBON-γ5β’2a/β’2mp/β’2mp_bilateral (MBON 01,03,04) and γd KCs promote sleep by activating MBON-γ2α?(MBON12)^33,56^. Our screen of the MB DAN cell types, revealed that PAM DANs projecting to lobe compartments containing the dendrites of wake-promoting MBONs are wake-promoting. We also found other subsets of PPL1 DANs that were wake-promoting and project to MB compartments that did not regulate sleep based on our screen of all KCs and MBONs^3^. Taken together, DANs projecting to sleep-regulating and neutral compartments to induce wakefulness but the mechanisms by which their activity is regulated and communicated to the core MB circuitry is unknown.

In addition to dopamine, GABA signaling also modulates sleep and wake microcircuits within MB^57^. The key source of GABA in the MB is anterior paired lateral neurons, APL and dorsal paired medial neurons DPM which are electrically coupled and increase sleep by GABAergic inhibition of wake-promoting α’/β’ KCs^57^. In the context of associative learning, there is strong evidence for interactions between KCs, APL and DANs^58,59^ but it is not clear if GABA and dopamine signaling represent opposing inputs to the KCs and MBONs in regulation of sleep.

Here we identify the precise circuit, cellular and molecular basis of how subsets of PAM DANs regulate wakefulness and test potential interactions between DA and GABA signaling. Specifically, we report that PAM DANs projecting to γ5, γ4 and β’2 MB compartments are required for sleep initiation and persistence and activity of PAM neurons is not directly modulated by sleep-need. Furthermore, we show that these effects are mediated by two of the four dopamine receptors, DopR1 and DopR2 that function within the KCs and MBONs of wake-regulating MB compartments.

The PAM DAN network with innervations exclusively in the MB represents a critical neuromodulator dependent sleep node that is distinct from CX. First, unlike dopamine mediated inhibition of sleep-promoting neurons within the dfb, the circuit we have identified is largely excitatory and mediates wakefulness by activating key wake-promoting regions (MBONs and KCs) of the MB. Second, the DANs projecting to CX function via DopR2 receptors, while in MB we find a role for both DopR1 and DopR2.

The PAM-DANs projecting to γ5, γ4 and β’2 MB compartments are particularly critical in understanding wakefulness as these neurons have been shown to be responsive to external stimuli such as sugar reward or electric shock used to reinforce memories in a cell-type specific manner^38,39,42^. Therefore, it’s likely that differential synaptic inputs and mode of communication to KC-MBONs via these DANs modulates MB function in promoting sleep, wakefulness or context dependent arousal.

## Materials and Methods

### Fly stocks and rearing conditions

All Fly stocks were maintained on cornmeal-agar-molasses medium (https://bdsc.indiana.edu/information/recipes/molassesfood.html) in 12 h light: 12 h dark conditions at 18°C with ambient humidity of 60–70%. The light intensity in the incubator was between 500-1200 lux measured using a luxmeter (Dr. Meter 1330B-V Digital Illuminance/Light Meter 0-200,000 Lux, Amazon Inc).

While, the fly stocks were maintained in the molasses media described above, virgin collection, genetic crosses and progeny collection for behavioral experiments were carried out in cornmeal dextrose agar media (https://bdsc.indiana.edu/information/recipes/dextrosefood.html). For all experiments 3-7-day old flies were collected and tested for behavioral experiments and immunohistochemistry.

All fly stocks were maintained and behavioral experiments were performed in Percival Scientific Inc. (DR36VL model) incubators. The following stocks used in the experiments were obtained from Bloomington Drosophila Resource Center:

26263 w [*]; P{y[+t7.7] w[+mC] =UAS-TrpA1(B). K} attP16
66600 y [1] w [*]; PBac {y[+mDint2] w[+mC] =20XUAS-TTS-shi[ts1]-p10} VK00005
31765 y [1] v [1]; P{y[+t7.7] v[+t1.8] =TRiP.HM04077} attP2
55239 y[1] sc[*] v[1]; P{y[+t7.7] v[+t1.8]=TRiP.HMC02344}attP2/TM3, Sb[1]
62193 y[1] sc[*] v[1]; P{y[+t7.7] v[+t1.8]=TRiP.HMC05200}attP40
26018 y[1] v[1]; P{y[+t7.7] v[+t1.8]=TRiP.JF02043}attP2
51423 y[1] sc[*] v[1]; P{y[+t7.7] v[+t1.8]=TRiP.HMC02893}attP2
65997 y[1] sc[*] v[1]; P{y[+t7.7] v[+t1.8]=TRiP.HMC06293}attP2
26001 y[1] v[1]; P{y[+t7.7] v[+t1.8]=TRiP.JF02025}attP2
36824 y[1] sc[*] v[1]; P{y[+t7.7] v[+t1.8]=TRiP.GL01057}attP2
50621 y[1] v[1]; P{y[+t7.7] v[+t1.8]=TRiP.HMC02988}attP40
31981 y[1] v[1]; P{y[+t7.7] v[+t1.8]=TRiP.JF03415}attP2
32194 w[*]; P{y[+t7.7] w[+mC]=20XUAS-IVS-mCD8::GFP}attP2
39171 w^1118^; P{GMR57C10-GAL4} attP2

Split-GAL4 lines: MB054B, MB312B, MB196B, MB194B, MB213B, MB060B, MB209B, MB060B, MB308B, MB438B, MB011B, MB010B, MB107B, MB298B, pBDGAL4, 58E02-GAL4, 48B04-GAL4 and 15A04-GAL4 were obtained from Dr. Yoshinori Aso and Dr. Gerry Rubin and have been described in^3,36,37,56^. For experiments using GAL4 lines, we used BDPGAL4U as a negative control, which contains the vector backbone used to generate each GAL4 line, but lacks any active enhancer motif to drive GAL4 expression^60,61^. Deletion mutants DopR1 (DopR1^attp^) and DopR2 (DopR2^attp^) were obtained from Todd Laverty and described in^62^. In both these deletion lines, the first coding exon has been deleted and replaced by an attP site. CaLexA flies (w*; P{LexAop-CD8-GFP-2A-CD8-GFP}2; P{UAS-mLexA-VP16-NFAT}H2, P{lexAop-rCD2-GFP}3/TM6B, Tb1) were obtained from Dr. Jing Wang and described in^63^.

### Sleep assays

For sleep experiments males and females were collected 3-7 days post-eclosion and placed in 65 mm × 5 mm transparent plastic tubes with standard cornmeal dextrose agar media, placed in a Drosophila Activity Monitoring system (Trikinetics), and locomotor activity data were collected in 1 min bins. Activity monitors were maintained in a 12 h:12 h light-dark cycle at 65% relative humidity, and tested flies were given 48 hours to acclimate to the light/dark cycle of the incubator. Total 24-h sleep quantity for each day of the experiment was extracted from locomotor activity data and sleep is defined as a contiguous period of inactivity lasting 5 min or more. Sleep profiles were generated depicting average sleep (minutes per 30 min) for the days of the experiment and maintained in the same tube. For CBZ experiments flies were placed on drug food the day prior to shibire inhibition as indicated in the experimental schematics in Figure 1, 2, S2 and S3.

**Figure 1:**
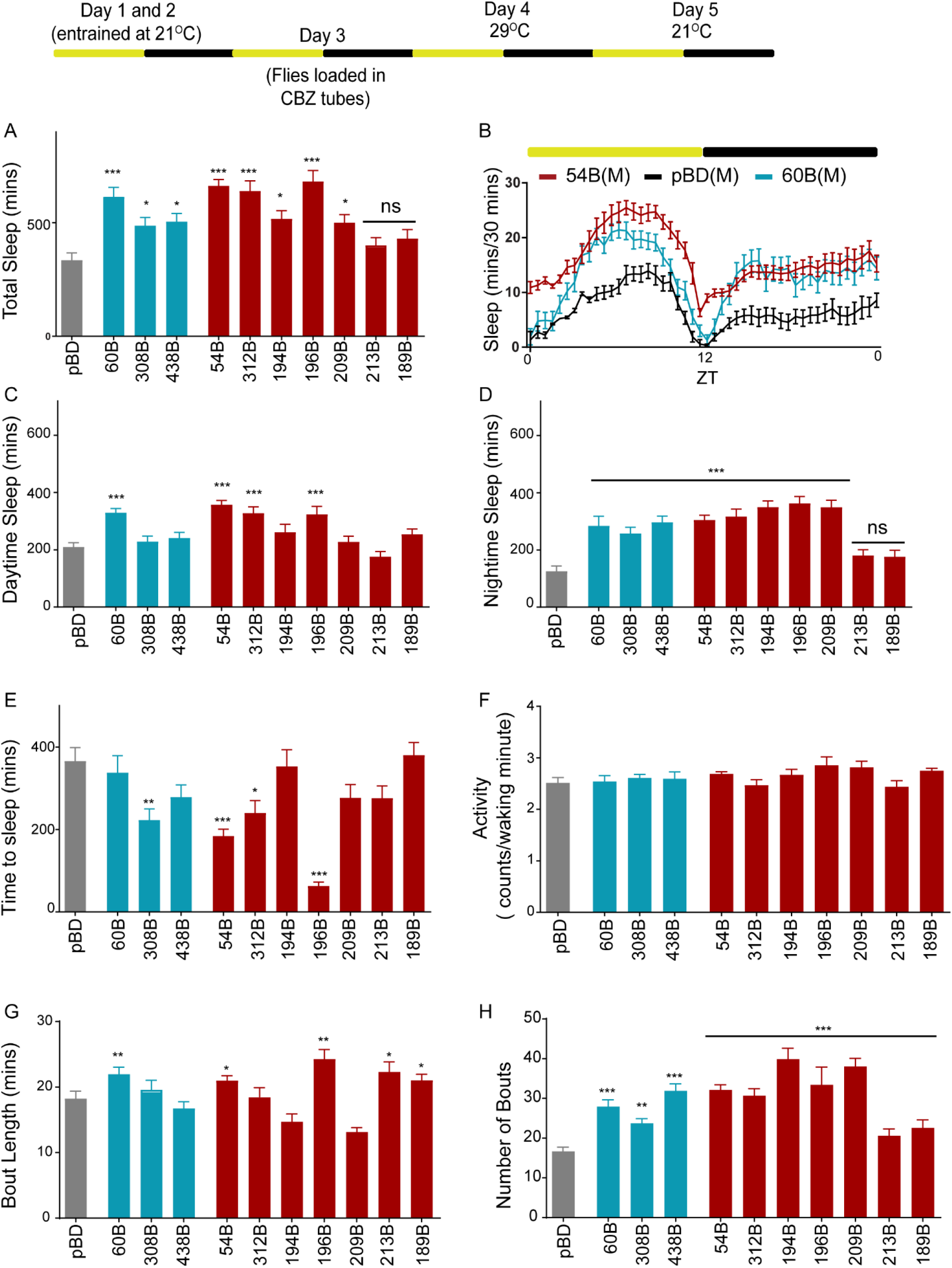
Wakefulness induced by pharmacologic suppression of GABA_A_ receptor, Rdl function requires subsets of MB dopamine neurons. Schematic of experimental protocol showing temperature and drug conditions over a 5-day period. Flies were entrained for Day 1 and 2 in vials. Males flies were loaded on 0.1mg/ml CBZ containing food (Day 3) in incubator maintained at 21°C. Day 4 the temperature was switched to 29°C and sleep was measured using the Drosophila Activity Monitoring system. (A) Total sleep in MB-DANs (PPL1-blue: MB060B, MB438B, MB308B and PAM-red: MB054B, MB312B, MB194B, MB196B, MB209B, MB213B and MB189B) where neural activity has been suppressed by over-expressing temperature sensitive dominant negative dynamin mutation, Shi^ts1^ in the presence of CBZ (Day 4). (B) Representative sleep profile of flies with targeted inhibition of specific PAM (MB054B, red, PPL1 (MB060B, blue) neurons with empty/enhancerless-GAL4 control (pBD, black). ZT indicates zeitgeber time where ZT 0: lights on and ZT 12: lights off. (C) Daytime sleep during the first 12-hour period during inhibition of specific PPL1 (blue), PAM (red) neurons as compared to empty-GAL4 control (pBD, grey). (D) Nighttime sleep during inhibition of specific PPL1 (blue), PAM (red) neurons as compared to empty-GAL4 control (pBD, grey). Statistical significance of total sleep, daytime and nighttime sleep was determined by performing one-way ANOVA followed by Dunnett post-hoc analysis [for Total Sleep, F (10, 912) =12.52, p<0.0001, for daytime sleep F (10, 912) =9.653, p<0.0001 and nighttime sleep, F (10, 912) =12.92]. For each of the experimental groups we had 73-96 flies which represented 4 independent experimental trials. Specifically, we had pBD (n=96), 54B (n=98), 60B (n=83), 189B (n=85), 194B (n=77), 209B (n=81), 213B (n=73), 308B (n=95), 312B (n=82), 438B (n=78) and 196B (n=75). (E) Sleep latency or time to sleep was calculated as the time gap between lights off and first sleep bout and analyzed using one-way ANOVA (F (10,912) =9.97, p<0.0001) followed by post-hoc Dunnett test. Latency [F (10,912) = 9.97, P<0.0001] was reduced in 54B, 312B and 196B which label PAM DANs and 308B that labels a PPL1 subset as compared to pBD. (F) Activity or average beam counts/waking minute were consistent between PAM, PPL1 and controls [F (10,921) =1.751, p<0.07] (G and H) Average bout length [H=90.25, p<0.0001] and bout number [H=163, p<0.0001] was analyzed by Kruskal Wallis ANOVA followed by Dunn’s post-hoc test. In figure panels A-H, ** indicates P<0.01 and *** indicates P<0.0001.

**Figure 2:**
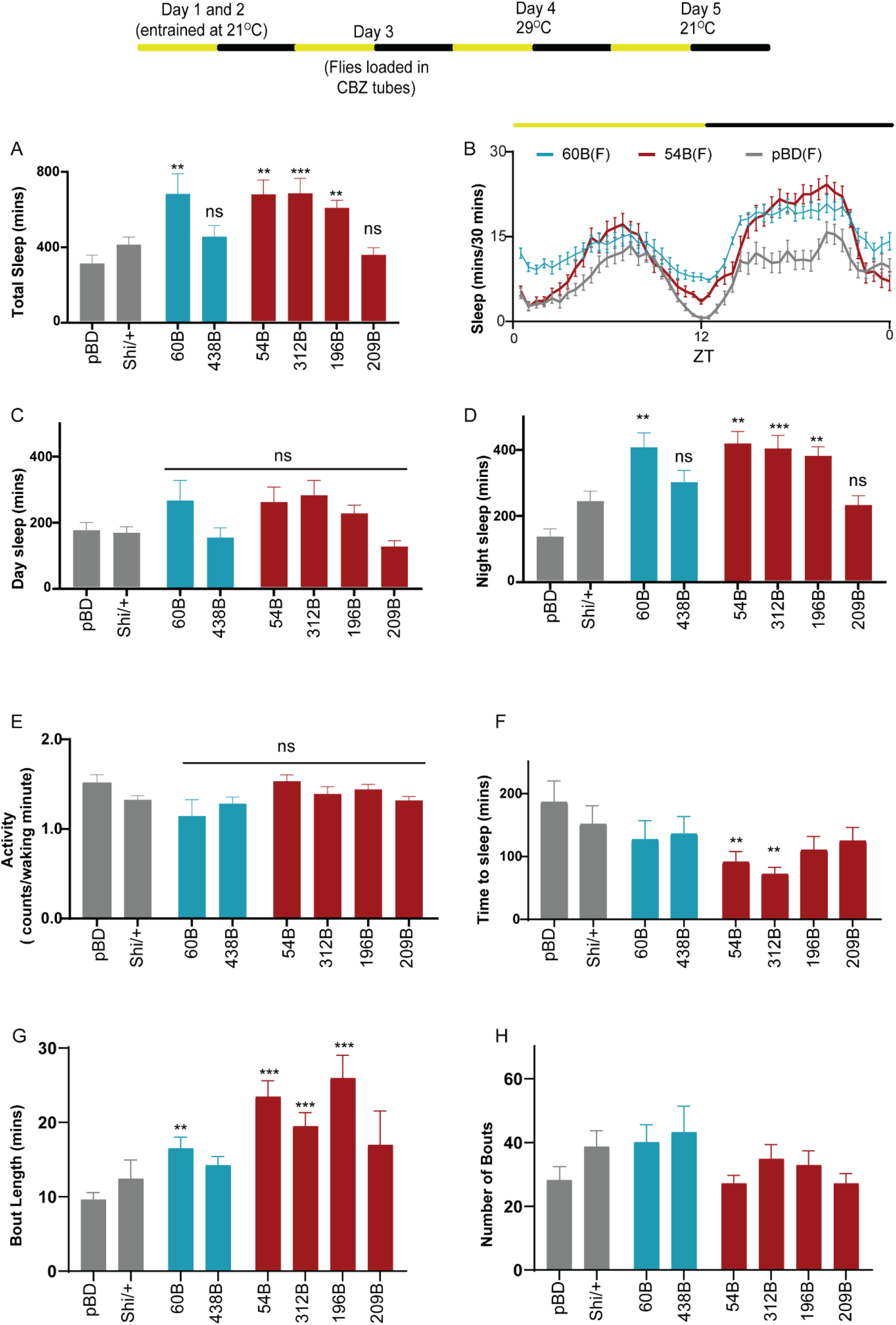
CBZ induced wakefulness requires dopamine signaling and is not sex specific. Schematic of experimental protocol showing temperature and drug conditions over a 5-day period. Flies were entrained for Day 1 and 2 in vials. Mated female flies were loaded on 0.1mg/ml CBZ containing food (Day 3) in incubator maintained at 21°C. Day 4 the temperature was switched to 29°C and sleep was measured using the Drosophila Activity Monitoring system. (A) Total sleep in MB-DANs (PPL1-blue: MB060B, MB438B and PAM-red: MB054B, MB312B, MB194B, MB196B, MB209B) where neural activity has been suppressed by over-expressing temperature sensitive dominant negative dynamin mutation, Shi^ts1^ in the presence of CBZ. (B) Representative sleep profile of flies with targeted inhibition of specific PAM (MB054B-black and MB312B-red) neurons with empty/enhancerless-GAL4 control (pBD, grey). ZT indicates zeitgeber time where ZT 0: lights on and ZT 12: lights off. (C) Daytime sleep during the first 12-hour period during inhibition of specific PPL1 (blue), PAM (red) neurons as compared to empty-GAL4 control (pBD, grey). (D) Nighttime sleep during inhibition of specific PPL1 (blue), PAM (red) neurons as compared to empty-GAL4 control (pBD, grey). Statistical significance of total sleep, daytime and nighttime sleep was determined by performing one-way ANOVA followed by Dunnett post-hoc analysis. For total Sleep [F (7, 238) =6.395, p< 0.0001], daytime sleep [F (7, 238) =3.010, p< 0.0048 and nighttime sleep [F (7, 238) =9.263, p< 0.0001]. For each of the experimental groups we had 28-36 flies which represented 2 independent experimental trials. Specifically, we had pBD (n=30), Shi/+ (n=30), MB060B (n=26), 209B (n=32), 312B (n=32), 438B (n=32), 196B (n=36) and 54B (n=28). (E) Activity or average beam counts/waking minute were consistent between PAM, PPL1 and controls [F (7,238) =1.474, p=0.17] (F) Sleep latency or time to sleep was calculated as the time gap between lights off and first sleep bout and analyzed using one-way ANOVA [F (7, 238) =2.352, p< 0.02] followed by post-hoc Dunnett test. Latency was reduced in 54B and 312B which label PAM DANs as compared to control. (G and H) Average bout length [H=6.633] and bout number [H=61.76] was analyzed by Kruskal Wallis ANOVA followed by Dunn’s post-hoc test. Bout length was elevated in all tested genotypes except MB438B and MB209B. In figure panels A-H, ** indicates P<0.01 and *** indicates P<0.0001.

All dTrpA1^64^ and Shibire^ts165^ experiments were conducted using temperature shift of 21°C and 29°C and RNAi experiments were conducted at 24°C in two Percival incubators (I 36VL). For RNAi experiments data represents an average of 2 days post-entrainment. For Shi and dTrpA1 experiments permissive temperature controls other genotypic controls were used for hit detection as indicated. Data analysis for sleep experiments was performed using MATLAB based software SCAMP developed by Dr. Christopher Vecsey (Skidmore College) and an earlier version of the software was published in^66^. For all screen hits, waking activity was calculated as the number of beam crossings/min when the fly was awake. Statistical comparisons between experimental and control genotypes were performed using Prism 7 (Graphpad Inc, CA).

Sleep deprivation was induced by intermittent shaking of the monitors displacing the flies at random intervals set by the DAM acquisition program as described in^33^. Sleep deprivation can be assessed by measuring sleep during deprivation in the sleep monitors as described above. After loading monitors, flies in the DAM monitor are placed on a horizontal vortexer. The vortexer power cord is attached to a light controller (LC4, Trikinetics Inc) that programs the power supply to the unit to deliver pulses that can be set using the DAM acquisition software. The pulse length and frequency can be controlled. We use 10 seconds of shaking per minute over a 16-hour period.

### Carbamazepine and Caffeine Feeding

CBZ (Sigma-Aldrich, C4024) was dissolved in 45% (2-hydroxypropyl)-beta-cyclodextrin (Sigma-Aldrich, H107) as described in to prepare a stock solution. For CBZ experiments, flies were loaded in tubes containing 2% agarose (Sigma, A9539) and 5% sucrose (S0389, Sigma-Aldrich) with 0.1 mg/ml CBZ. For caffeine feeding experiments Caffeine (Sigma-Aldrich, C0750) was mixed into melted sucrose/agar food at a concentration of 1mg/ml. Drug feeding experiments were conducted in sucrose-agarose media to allow for consistent mixing of the drug, which was difficult to monitor in the standard dextrose media because of its thicker consistency and opacity. All control (no drug) Shi^ts1^ experiments were conducted in sucrose/agarose media to account for sleep differences from different media.

### IHC

Dissection and immunohistochemistry of fly brains were performed as previously described with minor modifications (https://www.janelia.org/project-team/flylight/protocols). Brains of 3–7day old male flies were dissected in 1X PBS medium (BPP3920, Fisher Sci) and fixed in 2% paraformaldehyde (PFA, Electron Microscopy Sciences #15710) in PBT for 60 min at room temperature (RT). After washing in PBT (0.5% Triton X-100 from Sigma X100 in PBS), brains were blocked in 5% normal goat serum (NGS) (Vector Laboratories, S1000) in PBT overnight. Brains were then incubated in primary antibodies in NGS, nutated for 4 hours at room temperature, then transferred to 4°C for 2 days, washed three times in PBT for 30 min, then incubated in secondary antibodies diluted in NGS, nutated for 4 hours at room temperature, then transferred to 4°C for 2 days. Brains were washed thoroughly in PBT three times for 30 min and mounted in Vectashield (H-1000, Vector laboratories, CA) for imaging. The following antibodies were used: rabbit polyclonal anti-GFP (1:1000; Invitrogen), mouse anti-nc82 (1:50; Developmental Studies Hybridoma Bank, Univ. Iowa), and cross-adsorbed secondary antibodies to IgG (H+L): goat Alexa Fluor 488 anti-rabbit (1:800; Invitrogen) and goat Alexa Fluor 568 (1:400; Invitrogen).

For CaLexA experiment, images were taken under 40X magnification and analyzed using Fiji/ImageJ (https://imagej.net/Fiji) using a Nikon A1R resonant scanning confocal microscope. All the imaging conditions were kept constant within each experiment, except for Z-start and end settings and thickness of sample images was 100-120 microns.

To measure signal intensities, a maximal intensity projection of all the slices obtained from a z-stack comprising the PAM neuron (innervation patterns) was generated for control and experimental groups (sleep-deprived, carbamazepine and caffeine treated). ROI’s of projections in the MB region were manually drawn and identical in area for comparison groups. A second ROI in a region with no GFP expression was drawn and average pixel intensity was measures for all samples.

GFP signal was calculated as: Average pixel intensity of desired ROI-Average pixel intensity of background ROI. Comparison groups were scored single blind and GFP signal measured as fluorescence intensity was used for quantifying CaLexA signal.

### Statistical Analysis

Different sleep measures (sleep amount, latency, activity, bout length and number of bouts) are presented as bar graphs and represent mean ± SE or. A one-way ANOVA was used for comparisons between two or more treatments or two or more genotypes and post hoc analyses were performed using Dunnett’s correction. For data sets that did not follow a gaussian/normal distribution (bout numbers and bout length) we used non-parametric analysis (one-way ANOVA of ranks) and Kruskal Wallis Statistic. For comparisons of calcium levels as reported by CaLexA between genotypes or treatments, t test (two-tailed) was used. All statistical analyses and graphing were performed using Prism software (GraphPad Software 7.04; San Diego, California).

## Results

### 3.1 PAM and PPL1 DANs signaling is required for CBZ induced wakefulness

The 130 MB-DANs projects to and represent the primary modulatory input to MB and subdivided into PAM, PPL1 and PPL2 neurons. PAM neurons specifically project to the horizontal lobes (β, β’, and γ); while PPL1 neurons innervate the vertical lobes (α and α’), heel, and peduncle; and PPL2ab neurons project to the calyx^32^. In an unbiased screen of all DANs projecting to the MB using a cell specific split-GAL4 library we found that acute transient thermogenetic activation using the dTRPA1 temperature-gated depolarizing cation channel reveals multiple classes of PAM and PPL1 DANs suppresses sleep^3,33^. While, it’s not clear what modulates the activity of PAM and PPL1 DANs, GABA signaling suppresses sleep by APL neuron mediated inhibition of wake-regulating KCs^57^. Further, wake-promoting MBONs (MBON 01, 03, 04 and 05) are downstream of the wake-regulating KCs. Taken together, the KC-MBON network involved in wakefulness are regulated by PAM DANs and GABA signaling that regulates sleep antagonistically.

Carbamazepine or CBZ is a pharmacological agent that reduces GABAergic transmission by accelerating the desensitization of Rdl, Resistance to Dieldrin (GABA_A_ ionotropic receptor), and shown to suppress total sleep and increase sleep latency in a dose-dependent manner^67^. Further, *Rdl^MDRR^* mutants have enhanced GABAergic transmission due to altered channel properties of the Rdl receptors and exhibit shorter sleep latency and increased sleep^67^. While, the CBZ and Rdl effects on sleep are thought to be modulated by Pdf neurons^67,68^, the gene *Rdl* is expressed at high levels in the MB lobes and MBONs^41,69^. Genetic knock down of RDL enhances learning by increasing calcium influx into Kenyon cells/MB lobes that occurs when a fly is exposed to an odor^69,70^. Conversely, over-expression of Rdl in the MB lobes using broad GAL4 drivers increases GABA signaling and impairs aversive olfactory learning, where flies are conditioned to associate a neutral odor with electric shock^69,71^. Together, these data suggest GABA and Rdl signaling is prevalent in the MB.

In order to understand potential interactions between dopamine and GABA signaling, we inhibited synaptic output from sleep-regulating subsets of PAM and PPL1 neurons in the presence of CBZ, a known regulator of sleep and GABA transmission^57,67^. To investigate the role of compartment specific dopamine release by neurons we inhibited DANs by expressing temperature sensitive dominant negative mutation of dynamin, Shibire^ts1 65^. We blocked synaptic transmission in male and female flies from the sleep-regulating DANs in the presence and absence of 0.1mg/ml CBZ (carbamazepine) to test if DAN signaling is required for CBZ induced wakefulness.

We found that blocking dopamine release from smaller subsets of PPL1 neurons labelled by MB308B (PPL1 α3 and α’3), and MB438B (PPL1 MP1) increased total sleep both daytime and nighttime sleep, while a broader PPL1 driver MB060B (PPL1 α3, α’3, α’3α2 and γ2α’1) had stronger effects specific to daytime sleep. In comparison to the PPL1 neuronal inhibition, blocking dopamine release by specific PAM neurons (MB054B-PAM γ5, and MB312B-PAM γ4 and PAM γ4> γ1,2) resulted in much stronger suppression of the wake-inducing effects of CBZ (Figure 1A-D). Broader PAM lines (MB196B: PAM γ5, γ4, γ4>γ1,2, β’2a and MB194B: PAM γ5, β1, β’1, α1) had effects comparable to the broader PPL1 lines and MB213B (PAM β1 and β2) and MB189B (PAM γ5, β’2a, β’2m and β’2p) did not suppress wake induced by CBZ (Figure 1A-D). While, MB054B and MB189B label overlapping PAM γ5 neurons, they don’t affect sleep the same way. MB189B labels three β’2 subsets in addition to PAM γ5 and it is not clear if the three β’2 are wake-promoting or potentially mask the γ5 phenotype. While, several PAM γ5-GAL4 lines are wake-promoting when activated by dTrpA1, combinations of three β’2 subsets produced variable phenotypes^3^. Alternatively, MB189B could be a weaker driver as compared to MB054B that is not inhibited strongly by UAS-Shi^ts1^.

Taken together, we found that blocking dopamine release from both PAM and PPL1 neurons decreased CBZ induced latency or time to sleep after lights off at ZT 12. Like total sleep phenotype, latency was reduced strongly in multiple PAM drivers (MB054B, 312B and 196B) as compared to PPL1 (MB308B) (Figure 1E). Both PPL1 (MB060B) and PAM (MB054B and MB196B) inhibition increased bout length (Figure 1G) and number of bouts (Figure 1H) suggesting a role for these subsets in sleep initiation and maintenance. Activity measured as beam counts/waking minute were consistent between genotypes (Figure 1F) suggesting that the genetic manipulations and temperature elevation did not have differential effects on waking activity of the tested genotypes.

All Sleep measurements were conducted using the well-established infrared beam-crossing locomotor assay^72,73^ and flies were placed in food containing CBZ between ZT 3 to ZT 8 and sleep was recorded the day after at 21°C (permissive temperature-no inhibition) or 29°C (restrictive temperature-inhibition). Statistical significance of total sleep, daytime and nighttime sleep was determined by performing one-way ANOVA followed by Dunnett post-hoc analysis [for Total Sleep, F (10, 912) =12.52, p<0.0001, for daytime sleep F (10, 912) =9.653, p<0.0001 and nighttime sleep, F (10, 912) =12.92]. For each of the experimental groups we had 73-96 flies which represented 4 independent experimental trials. Sleep latency or time to sleep was calculated as the time gap between lights off and first sleep bout and analyzed using one-way ANOVA (F (10,912) =9.97, p<0.0001) followed by post-hoc Dunnett test. Latency [F (10,912) = 9.97, P<0.0001] was reduced in 54B, 312B and 196B which label PAM DANs and 308B that labels a PPL1 subset. Average bout length [H=90.25, p<0.0001] and bout number [H=163, p<0.0001] was analyzed by Kruskal Wallis ANOVA followed by Dunn’s post-hoc test.

The expression patterns with lobe specific innervation of these specific and broad sleep-regulating dopamine drivers were first described in^3,37^ and were confirmed by expression of GFP (Figure S1).

To test if genotypes tested in the study had differential sensitivity to CBZ we measured sleep in the presence of the drug at 21°C (permissive temperature). Flies expressing Shi^ts1^ in PAM and PPL1 neurons (60B, 308B, 438B, 54B, 312B, 189B, 196B, 194B, 209B and 213B) had reduced total sleep (~550-600 minutes, Figure S2A) [F (10, 469) =0.7056, p=0.7195], daytime sleep [F (10, 469) =0.8935, p=0.5391] Figure S2C, nighttime sleep [F (10, 469) =0.7018, p=0.7230] Figure S2D and increased latency [F (10, 469) =1.757, p=0.066], Figure S2E at permissive temperature and were not significantly different from control. Activity was also consistent between all tested genotypes F (10, 469) =1.240, p=0.2628, Figure S2F. These data (Figure S2) clearly show that inhibition of PAM/PPL1 at restrictive temperature results in increases in sleep in CBZ treated flies and is not a result of genotypic differences or differential sensitivity to CBZ between split-GAL4 lines (Figure S2).

To test if the effects of PAM and PPL1 DAN inhibition on CBZ induced wakefulness are sex-specific we tested mated females (Figure 2). We find that like males, inhibition of broad PPL1 driver MB060B and PAM driver lines MB196B, MB054B and MB312B suppresses CBZ induced wakefulness. Most of the sleep induced by DAN inhibition is observed during nighttime and caused by a combination of reduced latency (specifically in PAM lines MB054B and MB312B) and increased bout length (specifically in MB060B, MB054B, MB312B and MB196B). In flies expressing Shi^ts1^ in split-GAL4 lines (60B, 54B, 312B and 196B) total sleep (Figure 2A and B) was significantly higher as compared to control pBD [F (7, 238) =6.395, p< 0.0001]. Although, daytime sleep (Figure 2C) was elevated in PAM and PPL1 subsets [F (7, 238) =3.010, p< 0.0048], Dunnett’s post-hoc test did not reveal significant differences.

On the other hand, nighttime sleep (Figure 2D) was significantly elevated for PAM drivers (312B, 54B and 196B) and PPL1 drivers (60B and 438B) [F (7, 238) =9.263, p< 0.0001]. As in males, elevation in nighttime sleep was a combination of reduced latency [F (7, 238) =2.352, p< 0.02], Figure 2F in PAM drivers (312B and 54B) and increased bout length [H=61.76, p<0.0001], Figure 2G in PAM (312B, 196B and 54B) and PPL1 driver (60B). The number of bouts was similar between tested genotypes [H=6.633, ns], Figure 2H and activity was not significantly different between tested genotypes [F (7,238) =1.474, p=0.1770], Figure 2E. Taken together, in both males and females the wakefulness induced by CBZ and inhibition of GABA signaling is suppressed by blocking dopaminergic output from subsets of PAM (MB054B, MB196B and MB312B) and PPL1 neurons (MB060B).

Additionally, we also tested if the inhibition of subsets of PAM/PPL1 neurons in the absence of CBZ promotes sleep in both males and females (Figure S4). Acute thermogenetic activation of these subsets expressing dTrpA1 suppressed sleep with strong effects on nighttime sleep, when sleep is most consolidated^3^. However, we did not find significant difference in total sleep between flies at permissive (no inhibition, Day 1) and restrictive (inhibition, Day 2) temperature (Figure S4) in the absence of CBZ.

This could be a result of ceiling effects observed in nighttime sleep at permissive temperature or sleep loss occurring during nighttime as a result of temperature elevation to 29°C to block DAN output that makes it difficult to observe small increases during nighttime. Additionally, it’s highly likely that PAM/PPL1 DANs are inactive at nighttime and GABA sleep promoting effects are strong to allow the fly to sleep. Lastly, Shi^ts1^ is a weak regulator of neural activity^74^, Split-GAL4s are widely considered to be weaker than regular GAL4 lines^75^ and DAM measurement of sleep overestimates sleep amounts specifically during nightime^66^ adding to the challenges of observing increases in nighttime by inhibition of wake-promoting neurons^33^.

However, these results do not indicate that PAM-DANs are not required for natural wakefulness. Silencing of neuronal activity of various PAM DANs including MB196B and MB054B by expression of the hyperpolarizing kir2.1 K+ channel increases sleep^3^. We also find that activation of broad PAM GAL4 driver (76F05-GAL4) by dTrpA1 produces wakefulness specifically during nighttime, which is blocked by electrical silencing of subsets of PAM neurons targeted by 58E02-LexA (PAM γ5, γ4, γ4>γ1,2, β’2a) by Kir2.1 (an inward rectifying potassium channel) using the LexA-GAL4 system^3^.

In summary, PAM-DANs are required for natural wakefulness and CBZ induced arousal. While the effects of CBZ on sleep in flies are thought to be specific to GABAergic modulation of the Pdf neurons^67^, our data shows that CBZ effect on sleep is regulated in part by MB. This is supported by the abundance of Rdl receptors in KCs and their role in regulating calcium dynamics within MB lobes^69,70^. GABA and dopamine have known to work antagonistically within MB in regulating sleep and our data shows that CBZ produces wakefulness can be suppressed by blocking dopamine release to specific MB compartments PAM γ5, γ4 and β’2a.

To test the direct interactions between GABA signaling and activity of sleep-regulating PAM neurons we directly imaged activity in sleep-regulating PAM DANs MB196B and MB054B by expressing CaLexA^63^. We focused our attention on these neurons as PAM neurons had a stronger effect on suppressing sleep duration and latency effects induced by CBZ in both males and females and our work has previously identified downstream targets of these neurons allowing for a more detailed functional dissection of PAM mediated sleep regulation within the MB network^3,33,37,56^.

Like PAM-DANs (MB054B, MB312B and MB196B) synaptic silencing of the downstream wake-promoting MBONs 01,03, 04 and 05 (γ5, γ4 and β’2a) suppresses the wake-promoting effects of CBZ^3,33,37,56^. PAM DANs targeted by MB054B and MB196B specifically project to the γ5, β’2 and γ4 compartments of the mushroom body lobes and dendrites of wake-promoting MBONs tile these compartments.

To measure activity in PAM DANs in the presence of CBZ we used CaLexA or Calcium dependent nuclear import of LexA is a neural activity reporter that functions by Ca^++^ dependent nuclear import of a transcription factor which then drives GFP expression^63^. Hence, GFP levels in fixed tissue was used as a marker for neuronal activity to compare DAN activity in flies that were fed CBZ using the protocol identical to sleep recordings above. ROI’s of identical dimensions (area) were manually drawn to include the wake-regulating γ5, β’2 and γ4 MB (Figure 3) compartments and were compared in the presence and absence of CBZ which were dissected, stained and imaged in parallel using a single-blind protocol. MB054B projects to γ5 (Figure 3A and B), while, MB196B projects to γ5, β’2 and γ4 MB compartments (Figure 3A and C). We did not detect differences in calcium dependent GFP levels in sleep-regulating PAM DAN innervations in the absence and presence of CBZ for both MB054B and MB196B suggesting that exposure to the drug or reduced GABA transmission does not alter the directly activity of these neurons. The control and CBZ fed flies were compared using unpaired t-test with Welch’s correction (assumes unequal variances) for MB054B (t=0.021, df=18, p=0.9835, n=10) and MB196B (t=0.24, df=18, p=0.81, n=10).

**Figure 3:**
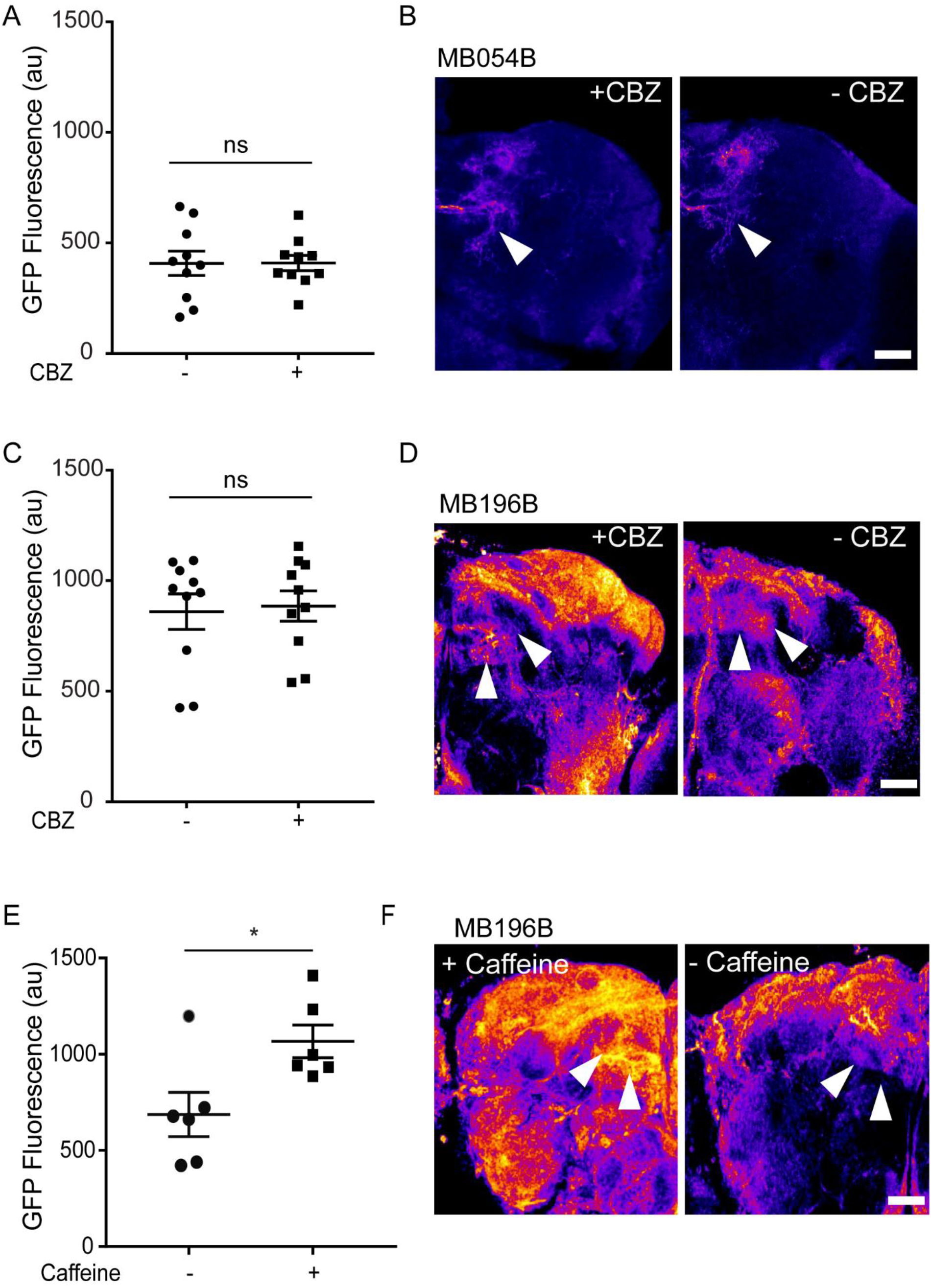
Pharmacological blocking of GABA transmission by CBZ does not directly modulate activity of PAM neurons. (A) MB054B projects to γ5 compartment of MB. We measured fluorescence signal in flies with PAM γ5 expressing CaLexA brains co-stained with antibodies against GFP and nc 82 between ZT 0-2 in flies that were treated with CBZ (~16-18 hours). CaLexA driven GFP+ fluorescence signal was quantified and compared between flies in CBZ and non-CBZ food. The control and CBZ fed flies were compared using unpaired t-test with Welch’s correction for MB054B (t=0.021, df=18, p=0.9835, n=10). (B) Maximum intensity projections of Z-stacks (8-10 slices) of two conditions (with and without CBZ) was pseudo colored with fire LUT to emphasize differential intensities. ROIs (arrowheads) were manually drawn to include MB compartments and were of equal area between groups. Scale bar represents 50um. (C) MB196B projects to γ5, γ4 and β’2 compartments of MB. We measured fluorescence signal in flies expressing CaLexA brains co-stained with antibodies against GFP and nc 82 between ZT 0-2 in flies that were treated with CBZ (~16-18 hours). CaLexA driven GFP+ fluorescence signal was quantified and compared between flies in CBZ and non-CBZ food. The control and CBZ fed flies were compared using unpaired t-test with Welch’s correction for MB196B (t=0.241, df=18 and p=0.81, n=10). (D) Maximum intensity projections of Z-stacks (10-15 slices) of two conditions (with and without CBZ) was pseudo colored with fire LUT to emphasize differential intensities. ROIs (arrowheads) were manually drawn to include MB compartments (γ5, γ4 and β’2) and were of equal area between groups. Scale bar represents 50um. (E) MB196B-GAL4-CalexA driven GFP+ fluorescence signal was quantified and compared between flies in caffeine (1mg/ml) and non-caffeine containing food to test the ability of CaLexA to report activity of DANs. The control and caffeine fed flies were compared using unpaired t-test with Welch’s correction (for MB196B t=2.66, df=9.25 and p=0.025, n=6). * indicates p<0.05. (F) Maximum intensity projections of Z-stacks (10-15 slices) of two conditions (with and without caffeine) was pseudo colored with fire LUT to emphasize differential intensities. ROIs (arrowheads) were manually drawn to include MB compartments (γ5, γ4 and β’2) and were of equal area between groups. Scale bar represents 50um.

The CaLexA system is well suited for visualizing neural activity in intact animals for long but not short time scales and used extensively in identifying sleep and wake-related activity within MB, CX (central complex) and clock neurons^27,76,77^. To ensure that the expression levels in split-GAL4 lines is strong enough to detect changes, we measured calcium dependent GFP levels in sleep-regulating PAM DAN’s in the absence and presence of caffeine, that was previously shown to affect sleep by modulating PAM activity. We found that 16-hour administration of 1mg/ml caffeine, increases activity of PAM neurons in MB196B which innervates to γ5, β’2 and γ4 MB compartments as previously published^78^. The control and caffeine fed flies were compared using unpaired t-test with Welch’s correction (for MB196B t=2.66, df=9.25 and p=0.025, n=6).

While the behavioral data show that PAM DAN activity to γ5, β’2 and γ4 MB compartments is required for wakefulness induced by reduced inhibitory GABA transmission, the calcium signaling or activity in PAM-DANs is not directly modulated by GABA. This suggests that either CBZ induced changes are small and undetectable by CaLexA or that GABA signaling affects sleep network at the level of KCs and MBONs that are downstream to PAM-DANs.

### 3.2 Activity of PAM DANs are not altered by sleep need

The PAM-DANs that produces the strongest sleep phenotype innervate the to γ5, β’2 and γ4 MB compartment and we recently showed that KCs and MBONs (MBON 01,03,04 and 05) within this compartment shows decreased electrical activity post sleep-deprivation when there is an increased sleep-need suggesting that neural activity within this compartment of the MB network is critical for sensing sleep-need and signaling wakefulness^33^. A recent study using EM dataset of a Full Adult Fly Brain (FAFB) mapped the inputs into the PAMγ5 DANs and identified that at least one PAM-γ5 DAN subtype (named γ5fb) receives recurrent synaptic input from γ5β’2a mushroom body output neurons (MBON 01,03 and 04)^79^

To test if the DANs projecting to wake-active γ5, β’2 and γ4 compartments of the MB alter their neural states as a function of sleep need like the downstream MBONs, we expressed CaLexA in these DANs (MB054B and MB196B) and measured neuronal activity post sleep-deprivation induced by 16-hour mechanical shaking as described in^33^. Sibling flies of the same genotype placed in the same temperature-controlled environment were used as sleep-replete controls (Figure 4A and B). There were no significant differences between sleep-deprived and sleep-replete conditions suggesting that unlike wake-promoting KCs and MBONs, PAM activity within γ5, β’2 and γ4 compartments is not altered by deprivation induced by sleep-need.

**Figure 4:**
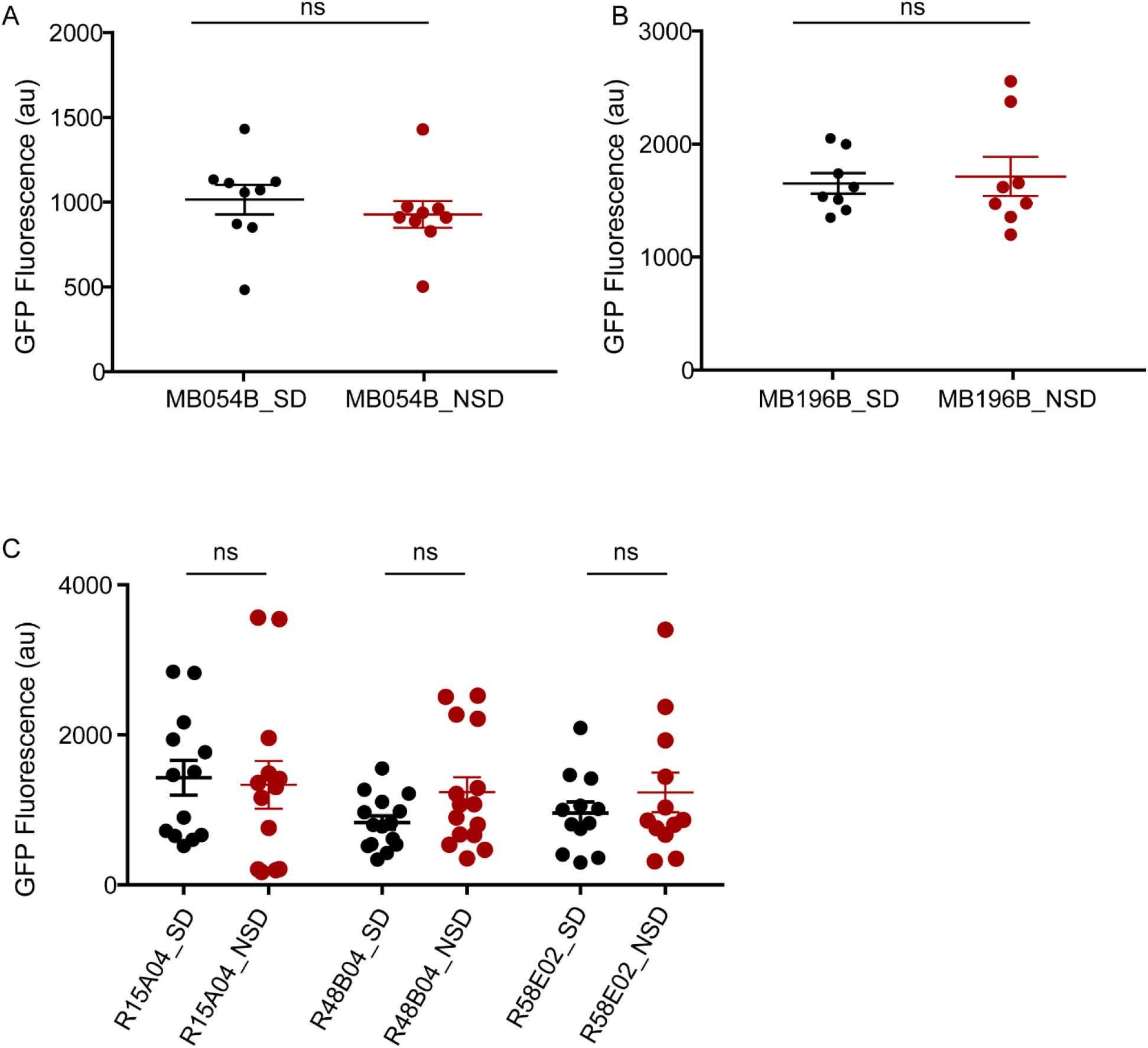
Unlike wake-regulating KCs and MBONs projecting to γ5, γ4 and β’2 MB compartments, sleep need does not alter activity and intracellular Ca^2+^ of upstream PAM neurons. (A and B) MB054B-GAL4 and MB196B-GAL4 expressing CaLexA brains were co-stained with antibodies against GFP and nc 82 between ZT 0-2 in flies that were sleep-deprived (for 16 hours) or sleep replete. CaLexA driven GFP+ fluorescence signal was quantified and compared between sleep-deprived and sleep replete controls. CaLexA signal was compared between sleep-deprived and sleep-replete groups using unpaired t-test with Welch’s correction for MB054B (t=0.75, df=16 and p=0.46, n=9) and MB196B (t=0.68, df=14 and p=0.504, n=8). (C) Broader PAM-GAL4 drivers, 15A04-GAL4, 48B04-GAL4 and 58E02-GAL4 lines were used to drive CaLexA expression and GFP fluorescence was compared between sleep-deprived and sleep-replete states in fly brains between ZT 0-2. CaLexA signal was compared between sleep-deprived and sleep-replete groups using unpaired t-test with Welch’s correction for 15A04-GAL4 (t=0.24, df=24 and p=0.81, n=13), 48B04-GAL4 (t=1.8, df=28 and p=0.08, n=15) and 58E02-GAL4 (t=0.91, df=22 and p=0.38, n=12).

CaLexA signal was compared between sleep-deprived and sleep-replete groups using unpaired t-test with Welch’s correction for MB054B (t=0.75, df=16 and p=0.46, n=9) and MB196B (t=0.68, df=14 and p=0.504, n=8). Since, GFP was visible in both conditions it is possible that there is basal activity in these neurons that does not change with sleep-deprivation. Additionally, the changes in activity could be very small or undetectable because split-GAL4 lines are weaker drivers of UAS-transgenes.

To address this possibility of expression strength of split-GAL4 lines, we used three additional GAL4 lines (15A04-, 48B04- and 58E02-) that label these sleep-regulating PAM DANs. Like split-GAL4 lines, some basal activity was detected in both sleep-deprived and sleep-replete flies but were not significantly different (Figure 4C). CaLexA signal was compared between sleep-deprived and sleep-replete groups using unpaired t-test with Welch’s correction for 15A04-GAL4 (t=0.24, df=24 and p=0.81, n=13), 48B04-GAL4(t=1.8, df=28 and p=0.08, n=15) and 58E02-GAL4 (t=0.91, df=22 and p=0.38, n=12).

Together these data suggest that while dopamine release regulates sleep and wakefulness, the activity of these neurons is not directly regulated by GABA transmission and homeostatic mechanisms. These findings are consistent with previously published results where TrpA1-based stimulation of sleep-regulating PAM neurons (MB054B, MB312B, MB196B) caused a rapid and robust decrease in nighttime sleep, but were not followed by a positive sleep rebound the day after stimulation^3^. Since, PAM-DAN activity has to be closely matched by receptivity of Dopamine signal within the downstream KCs and MBONs that are influenced by GABA signaling and sleep-need we focussed on exploring the molecular nature of DA receptors involved in PAM-DAN network.

### 3.3 Sleep regulating DANs signal via DopR1 and DopR2

Four dopamine receptors (all G-protein coupled receptors) have been identified in the Drosophila genome: DopR1, DopR2, D2R and DopEcR^80-83^. As in humans, DopR1 and DopR2 are D1-like receptors and functions via activation of the cAMP pathway, while D2-like receptors inhibit this pathway. Hence, the effect of DA on a specific postsynaptic neuron depends on the type of DA receptor that is expressed. Dopamine receptors DopR1 and DopR2 are highly expressed in the MB (KCs and MBONs) and have been shown to increase production of cAMP through in vitro assays^38,81,84,85^. However, mutant studies highlight a dual, opposing role for these receptors in regulating memory, in which DopR1 acts to promote memory formation while DopR2 serves to degrade memory^86,87^.

Further, loss-of-function mutations of D1 dopamine receptor *DopR* are shown to enhance repetitive air puff startle-induced arousal (Resh), while increasing nighttime sleep. Expression and restoration of DopR in the mutant background specifically in the central complex but not the MB rescues the Resh phenotype suggesting that arousal and wakefulness can be disassociated in the context of Resh and that *DopR* functions differentially within distinct neural structures in regulation of wakefulness and stimuli dependent arousal^88^. Several DA receptor or transporter mutations have been shown to decrease arousal thresholds (to air puffs, light or mechanical stimuli) in awake flies, independent of their role in sleep^25,89,90^. Hence, both the compartmentalization of DA clusters in the fly brain and distinct post-synaptic effects exerted by different receptors within multiple neural substrates underlies the complex role of dopamine in regulating endogenous arousal (wakefulness) and exogenous arousal (behavioral responsiveness to sensory stimuli).

While, the split-GAL4 based neuronal targeting helps identify the specific sources of DA involved in endogenous arousal behaviors like wakefulness, the post-synaptic effects are more complex to pin down. Sleep-regulating PAM DANs innervating the γ5, β’2 and γ4 MB compartments which is composed of KCs and MBONs that have reduced electrical activity post sleep-deprivation^33,91^.

Two of the dopamine receptors, DopR1 and DopR2 are co-expressed at high levels in the specific populations of Kenyon Cells (KCs)and MBONs that form the γ5, β’2 and γ4 MB compartments^86,87,92,93^. While, the role of the DopR2 in sleep regulation and inhibition of dfb neurons within central complex is well characterized, the identity of dopamine receptors involved in MB specific sleep regulation is unknown^23,24^.

We used a pan-neuronal driver nsyb-GAL4 (R57C10) with dicer expression and targeted all four dopamine receptors using validated UAS-RNAi lines in the context of sleep regulation. Given the wide variety of RNAi of lines available to downregulate receptor transcripts we picked at least two transgenic lines for each receptor that have been previously validated by quantitative RT PCR^40,41,51,58,94^. Male flies with reduced receptor levels were tested and we found that downregulation of DopR1 (two RNAi lines: 31765 and 62193) and DopR2 (one RNAi line: 65997) specifically increased total sleep without altering the locomotor activity measured by beam counts/minute during wake-period (Figure 5A and 5D). Both these receptors had differential effects on daytime and nighttime sleep such that Dop1R1 affected both daytime and nighttime sleep while DopR2 effects were restricted to nighttime sleep (Figure 5B and 5C). Consistent with blocking DA release, we found that one of the DopR1 and DopR2 RNAi lines (31765 and 65997) reduced sleep latency. Bout length and number of bouts were consistent between tested genotypes with the exception of one RNAi line targeting DopR2. While, all of these RNAi lines have been previously validated to reduce receptor expression, it’s not clear if the downregulation is variable or if the receptors have redundant functions. Further, ceiling effects in nighttime sleep measurements potentially undermines the small increases in sleep and leads to false negatives even when the RNAi lines are effective in downregulating receptor levels. Taken together, pan-neuronal knockdown of DopR1 and DopR2 increased sleep and decreased sleep latency consistent with the role of DAN signaling in the MB.

**Figure 5:**
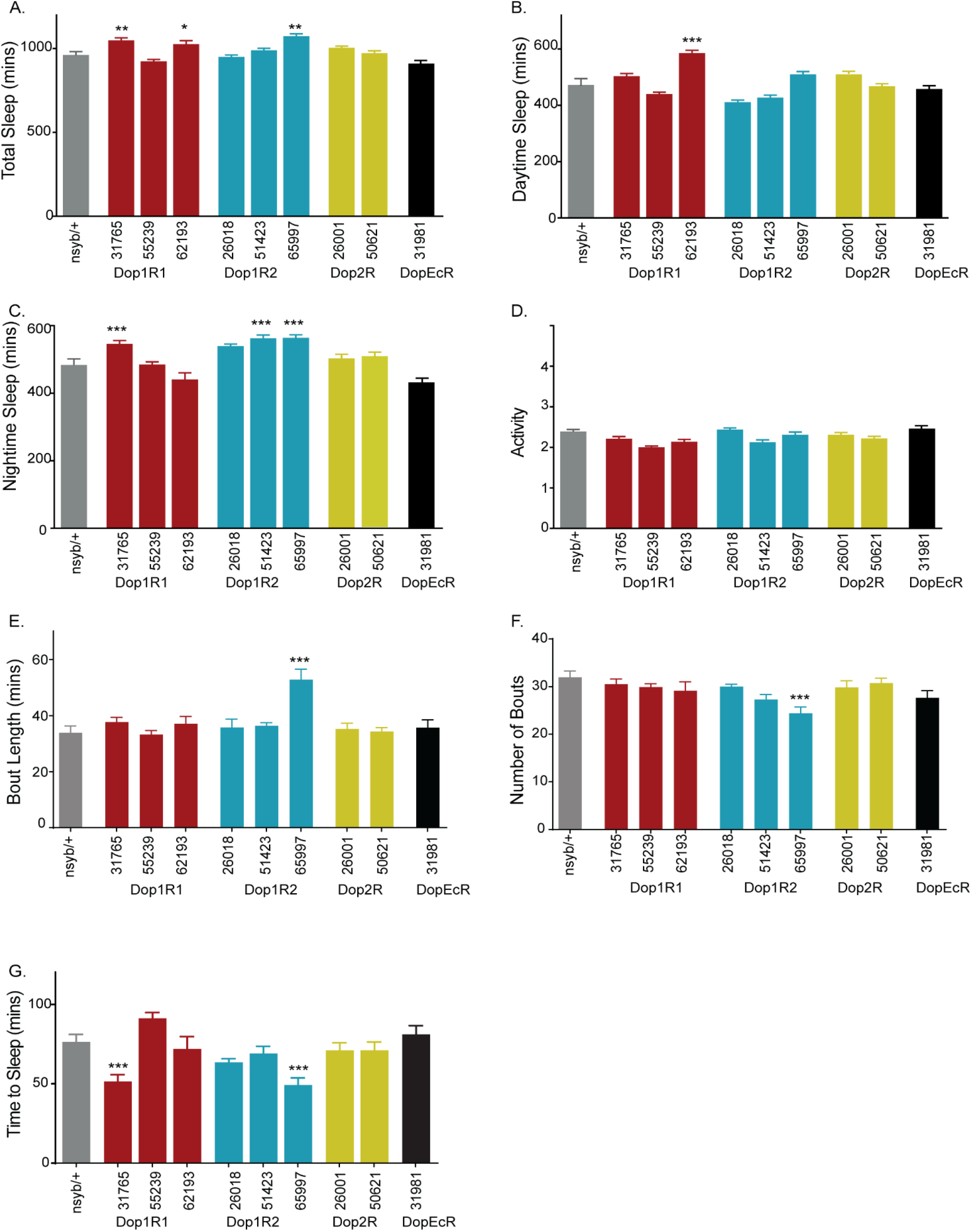
Pan-neuronal knockdown of dopamine receptors by validated RNAi lines increases sleep amount without altering waking activity. (A) Total sleep in male flies expressing validated UAS-RNAi targeting DopR1 (3 lines: 31765, 55239 and 62193-red), DopR2 (26018,51423 and 65997-blue), Dop2R (26001 and 50621-yellow) and DopEcR (31981-black) pan-neuronally using nsyb-GAL4. Sleep data represents 2-day average (Day 3 and 4) at 24°C after 2-day entrainment. nsyb-GAL4/+ (grey bars) was used as negative control. (B and C) Daytime and nighttime sleep of nine DA receptor RNAi lines and control. Statistical significance of total sleep, daytime and nighttime sleep was determined by performing one-way ANOVA followed by Dunnett’s post-hoc analysis [for Total Sleep= F (9, 938) = 13.64, p<0.0001, for daytime sleep F (9, 938) = 28.45, p<0.0001 and for nighttime sleep F (9,938) = 15.65, p<0.0001]. For each of the experimental groups tested we had 88-99 flies which represented 4 independent experimental trials, sample included nsyb/+ (95), nsyb/31765 (96), nsyb/55239 (92), nsyb/62193(98), nsyb/26018(96), nsyb/51423(98), nsyb/65997 (93), nsyb/26001 (96), nsyb/50621(96) and nsyb/31981(88). (D) Activity or average beam counts/waking minute were consistent between tested genotypes and controls [F (9,938) =30.50]. (E and F) Average bout length [H=50.54] and bout number [H=30.61] was analyzed by Kruskal Wallis ANOVA followed by Dunn’s post-hoc test. Bout length was elevated in nsyb/DopR2-RNAi (65997). (G) Sleep latency or time to sleep was calculated as the time gap between lights off and first sleep bout and analyzed using one-way ANOVA [F (9, 939) =9.505, p<0.0001] followed by post-hoc Dunnett test. Latency was reduced in nsyb/DopR1-RNAi (31765) and nsyb/DopR2-RNAi 65997) flies as compared to nsyb/+.

Statistical significance of total sleep, daytime and nighttime sleep was determined by performing one-way ANOVA followed by Dunnett’s post-hoc analysis [for Total Sleep= F (9, 938) = 13.64, p<0.0001, for daytime sleep F (9, 938) = 28.45 and for nighttime sleep F (9,938) = 15.65]. For waking activity, beam counts/ waking was consistent between genotypes F (9,938) =30.50 and time to sleep after lights off (sleep latency) was reduced in two genotypes F (9,938) =9.505, p<0.0001. Sleep bout length (H=50.54) and bout number (H=30.61) was not different between groups.

### 3.4 DopR1 and DopR2 regulate sleep amount, bout characteristics and sleep latency by influencing specific MB compartments

Since, pan-neuronal manipulations affect receptor levels outside of MB we repeated these experiments with validated RNAi lines that target DopR1 (31765 and 62193) and DopR2 (65997) that increased sleep (Figure 5, DopR1). We targeting DopR1 and DopR2 knockdown to MB neuronal populations that are potentially downstream to the to wake-active γ5, β’2 and γ4 PAM DANs using highly specific split-GAL4 lines described in^37,56^. All the RNAi lines used were inserted in the same genomic location on the 3^rd^ chromosome for comparable expression. Specifically, we targeted two MB output neurons (MBONs) projecting to the γ5 (MB011B), β’2 (MB011B) and γ4 (MB298B) synaptic compartments and Kenyon cell populations projecting to these lobes (MB010B-all KCs, MB107B-α’ß’ KCs).

One of the RNAi lines targeting DopR1 transcripts (31765) increased total sleep when expressed in γ5 β’2 (amp) MBONs (MBON 01, 03 and 04), α’ß’ KCs and all KCs but not in γ4 MBONs as compared to pBD (control) suggesting that suppression of dopamine signaling via this receptor subtype increases total sleep [F (4, 196) = 18.79, p<0.0001] in MB011B, MB107B and MB010B (Figure 6A). Knockdown of DopR1 in MB011B (γ5 β’2 (amp)) increases daytime sleep but reduced daytime sleep in MB0298B (γ4) ([F (4, 196) = 37.21, p<0.0001] (Figure 6C). Nightime sleep [F (4, 196) = 42.36, p<0.0001] in all downstream neurons (MB011B, MB10B, MB107B and MB 298B) was elevated as compared to control (Figure 6D). The most drastic behavioral effect however was observed in reduced latency (Figure 6E) phenocopying the effects of reduced DA signaling.

**Figure 6:**
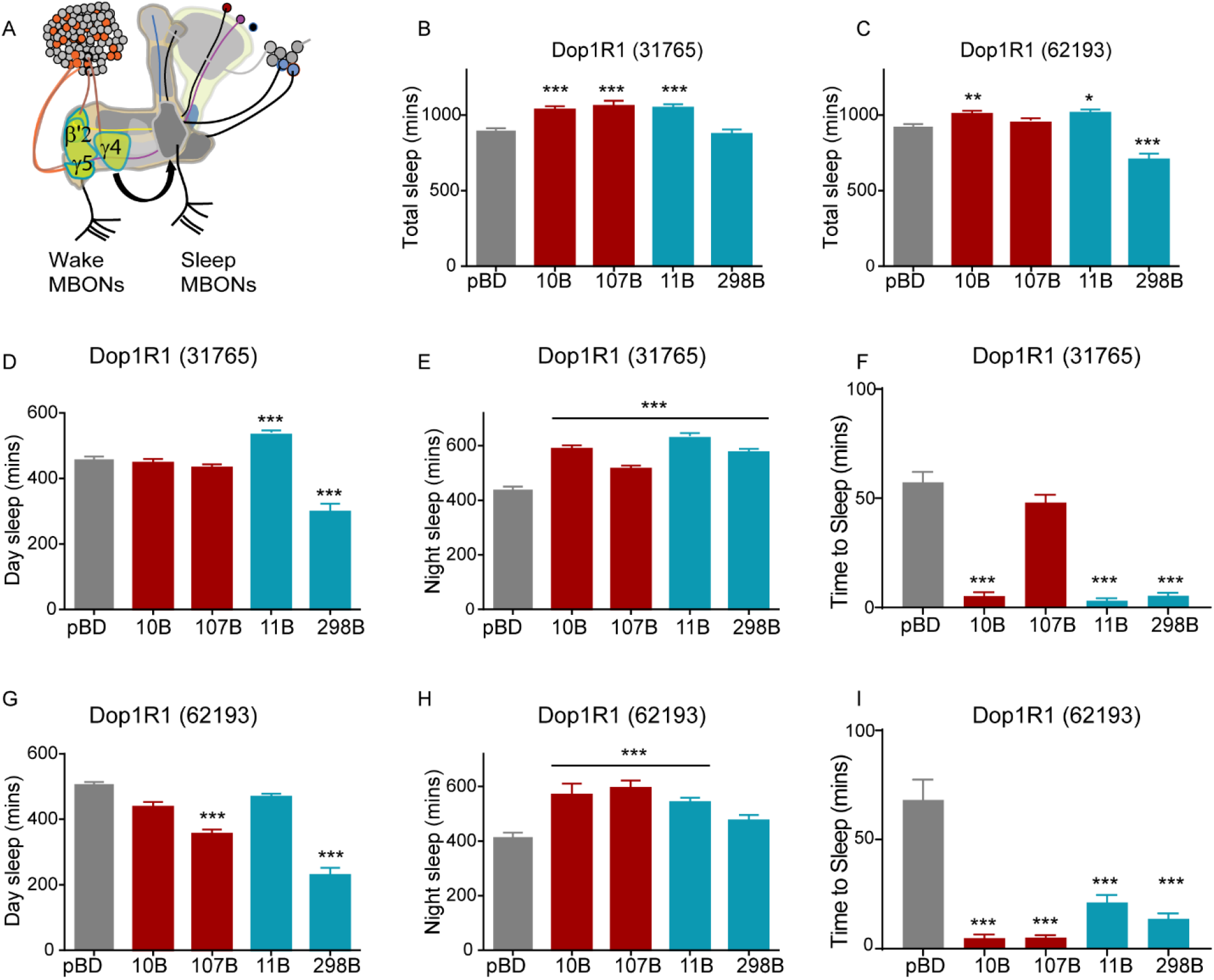
Knockdown of DopR1 MB-KCs and MBONs of the sleep-regulating γ5 and β’2 compartment increased sleep. (A) Schematic of MB sleep-regulating γ5, γ4 and β’2 compartments that represents the interactions between KCs, MBONs and PAM-DANs. (B) Total Sleep during a 24-hour period in flies expressing UAS-DopR1 RNAi (31765) in MB-KCs (MB010B, MB107B), MBONS (MB011B, MB 298B) and pBD. One-way ANOVA [F (4, 196) = 18.79, p<0.0001], post-hoc Dunnett’s test reveals increased sleep in MB010B, MB107B, MB011B but not MB 298B. For each of the experimental groups tested we had 37-48 flies which represented 2 independent experimental trials, sample included 31765/pBD (48), 31765/MB010B (40), 31765/MB107B (38),31765/MB011B (38),31765/MB298B (37). (C) Total Sleep during a 24-hour period in flies expressing UAS-DopR1 RNAi (62193) in MB-KCs (MB010B, MB107B), MBONS (MB011B, MB 298B) and empty-GAL4 (pBD). One-way ANOVA [F (4, 220) = 23.44, p<0.0001], post-hoc Dunnett’s test reveals increased sleep in MB010B and MB011B but not in MB107B and MB 298B. For each of the experimental groups tested we had 39-47 flies which represented 2 independent experimental trials, sample included 62193/pBD (47), 62193/MB010B (39), 62193/MB107B (44), 62193/MB011B (43), 62193/MB298B (47). (D and E) Daytime and Nighttime Sleep in flies expressing UAS-DopR1 RNAi (31765) in MB-KCs (MB010B, MB107B), MBONS (MB-11B, MB 298B) and empty-GAL4 (pBD). One-way ANOVA for daytime [F (4, 196) = 37.21, p<0.0001] and nighttime [F (4, 196) = 42.36, p<0.0001] followed by post-hoc Dunnett’s test reveals increased daytime sleep in MB011B and reduced daytime sleep in MB298B. Nighttime sleep was significantly different in all tested genotypes as compared to pBD control. (F) Sleep latency or time to sleep (minutes) was calculated as the time gap between lights off and initiation of the first sleep bout for flies expressing UAS-DopR1 RNAi (31765) in MB-KCs (MB010B, MB107B), MBONS (MB-11B, MB 298B) and empty-GAL4 (pBD). One-way ANOVA for latency [F (4, 196) = 42.08, p<0.0001] followed by post-hoc Dunnett’s test reveals significantly reduced latency in MB010B, MB011B and MB298B but not MB107B. (G and H) Daytime and Nighttime Sleep in flies expressing UAS-DopR1 RNAi (62193) in MB-KCs (MB010B, MB107B), MBONS (MB-11B, MB 298B) and empty-GAL4 (pBD). One-way ANOVA for daytime [F (4, 220) = 66.61, p<0.0001] and nighttime [F (4, 220) = 45.31, p<0.0001] followed by post-hoc Dunnett’s test reveals daytime sleep is reduced in MB010b, MB107B and MB298B. Nighttime sleep was significantly different in all tested genotypes as compared to pBD control. (I) Sleep latency or time to sleep (minutes) was calculated as the time gap between lights off and initiation of the first sleep bout for flies expressing UAS-DopR1 RNAi (62193) in MB-KCs (MB010B, MB107B), MBONS (MB-11B, MB 298B) and empty-GAL4 (pBD). One-way ANOVA for latency [F (4, 220) = 26.84, p<0.0001] followed by post-hoc Dunnett’s test reveals significantly reduced latency in MB010B, MB107B, MB011B and MB298B.

A closer analysis of the sleep structure reveals that latency was not the only driver of increased nighttime sleep and that average bout length (H=50.28, Figure S4A) was higher in MB010B, MB107B and MB011B. While, the number of bouts (H=32.97, Figure S4B) were reduced in MB107B but not the other cell-types. Taken together, these data show that knockdown of DopR1 receptor affects total sleep by reducing latency and increasing the duration of average sleep bout in KCs and MBONs. However, the increased sleep phenotype in α’ß’ KCs was mostly contributed by increased sleep bout length and decreased number of bouts. Activity was consistent between tested genotypes [F (4,196) =3.424], Figure S4E and that modulating receptor levels did not affect activity during wake bouts consistent with previous findings of DopR manipulations in MB^95^.

A second RNAi line targeting the same receptor (DopR1) increased total sleep when expressed in all KCs (MB010B) and γ5 β’2 (amp) MBONs (MB011B) [F (4, 220) = 23.44, p<0.0001] without altering total sleep in α’ß’ KCs (MB107B) and reducing sleep in γ4 MBONs (MB298B) (Figure 6B). Daytime sleep [F (4,220) =66.61, p<0.0001) was more variable with the knockdown of this RNAi line and all tested lines except MB011B had reduced sleep as compared to empty split-GAL4 control. Sleep duration during nighttime when sleep is more consolidated was elevated in all the tested genotypes, which is the likely contributor of observed increases in total sleep [F (4,220= 45.31, p<0.0001)]. Further, we find that the increases in nighttime sleep resulted from decreased sleep latencies [F (4,220) = 26.84, p<0.0001) in flies where the DopR1 levels were downregulated specifically in all Kenyon cells and γ5 β’2 (amp) MBONs. Like, the first DopR1 RNAi line (31765), increase in nighttime sleep was also contributed by increase in length of average sleep bout (H=48.56) in MB107B and MB011B. Sleep bout number (H=49.70) was mostly consistent between genotypes expect for MB010B which labels all KCs.

In summary, two transgenes encoding RNAi lines targeting DopR1 showed consistent increase in total sleep, decrease in sleep latency and increased sleep bout length when expressed in MBON 01-04, α’ß’ KCs and all KCs. However, the effects on γ4 MBONs (MB298B) are perplexing as it reduces or has no effect on total sleep specifically during daytime, while decreasing sleep latency. The PAM-DANs projecting to γ4 MBON are labelled by MB312B and are strongly wake promoting and this is consistent with experiments where activation of γ4 MBON also induce wakefulness^56^. It is likely that MB312B releases a co-transmitter that influences wakefulness via γ4 MBON in a dopamine independent way. Recent evidence shows that nitric oxide synthase, NOS an enzyme required for synthesis of NO has been detected in several DANs and works antagonistically to dopamine in regulating olfactory memory^47^. Whether, such a co-transmitter dependent mechanism operates within MB in sleep regulation is unclear but could be a potential mechanism in diversifying neural signaling and supporting multiple outcomes in regulation of endogenous (wakefulness) and exogenous arousal.

Using, the above cell type specific regulation of DA receptors we downregulated the second D1 receptor, DopR2 receptor function (65997) in wake-regulating MBON compartments and Kenyon cell populations. We found that reduction in DA signaling via DopR2 receptor increased total sleep [F (4,228=46.49, p<0.0001] when expressed in γ5 (MB011B), β’2 (MB011B) and Kenyon cell populations projecting to these lobes (MB010B-all KCs, MB107B-α’ß’ KCs) but not in γ4 (MB298B) MBONs (Figure 7A). This differential effect on MB011B and MB298B was similar to that observed for DopR1 (Figure 6).

**Figure 7:**
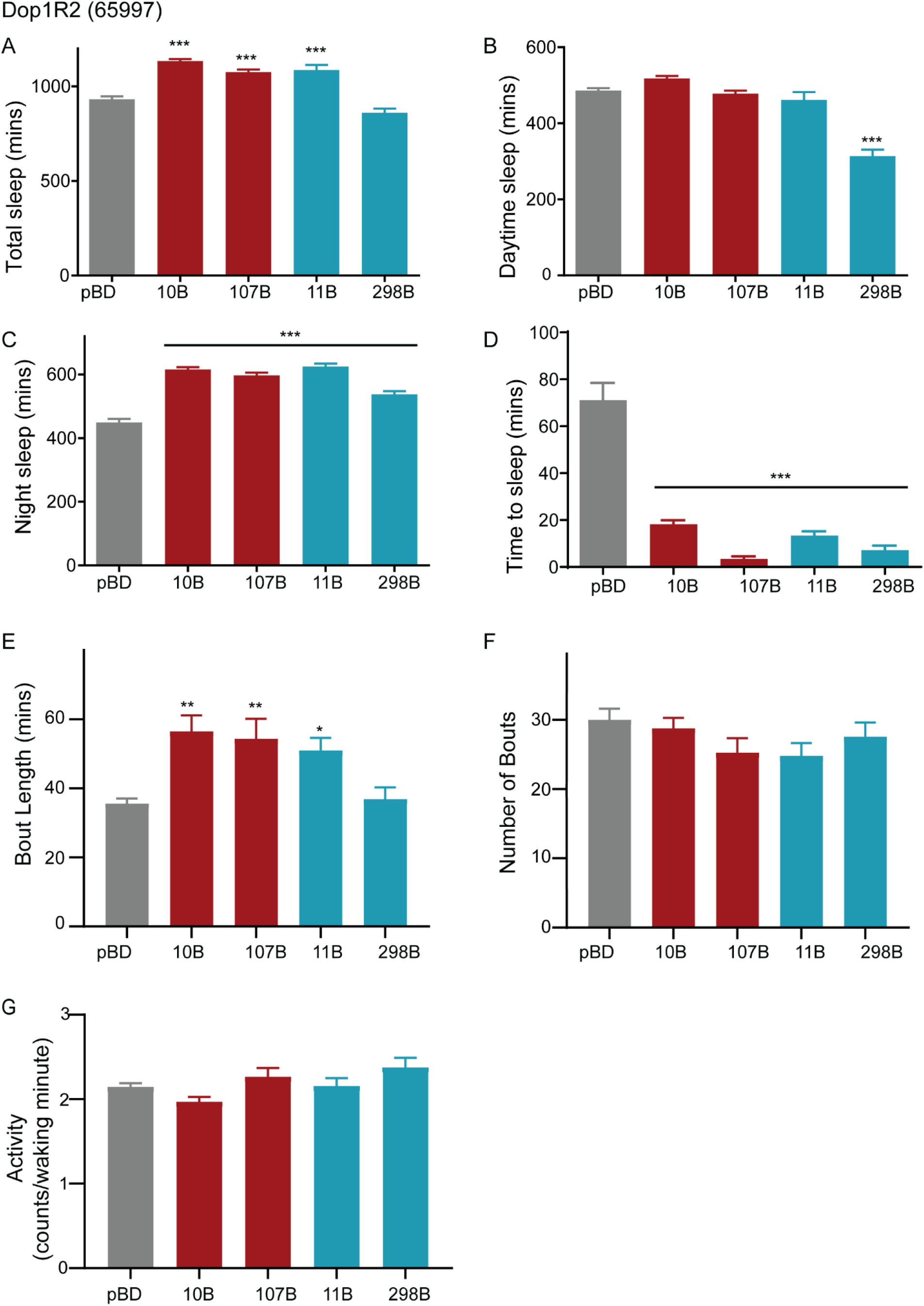
Knockdown of DopR2 in sleep-regulating MB-KCs and MBONs increased sleep and reduced latency. (A) Total Sleep during a 24-hour period in flies expressing UAS-DopR2 RNAi (65997) in MB-KCs (MB010B, MB107B), MBONs (MB-11B, MB 298B) and empty-GAL4 (pBD). One-way ANOVA [F (4, 228) = 46.49, p<0.0001], post-hoc Dunnett’s test reveals increased sleep in MB010B, MB107B, MB011B but not MB 298B. For each of the experimental groups tested we had 44-48 flies which represented 2 independent experimental trials, sample included 65997/pBD (48), 65997/MB010B (46), 65997/MB107B (47), 65997/MB011B (44), 65997/MB298B (48). (B and C) Daytime and Nighttime Sleep in flies expressing UAS-DopR2 RNAi (65997) in MB-KCs (MB010B, MB107B), MBONS (MB-11B, MB 298B) and empty-GAL4 (pBD). One-way ANOVA for daytime sleep [F (4, 228) = 65.45, p<0.0001] and nighttime sleep [F (4, 228) = 86.04, p<0.0001], post-hoc Dunnett’s test reveals increased nighttime sleep in MB010B, MB107B, MB011B and decreased daytime sleep as compared to control. (D) Sleep latency or time to sleep (minutes) was calculated as the time gap between lights off and initiation of the first sleep bout for flies expressing UAS-DopR2 RNAi (65997) in MB-KCs (MB010B, MB107B), MBONS (MB011B, MB 298B) and control (pBD). One-way ANOVA for daytime sleep [F (4, 228) = 40.17, p<0.0001] followed by post-hoc Dunnett’s test reveals significant decrease in latency in all tested genotypes as compared to control. (E and F) Average bout length and bout number in flies expressing UAS-DopR2 RNAi (65997) in MB-KCs (MB010B, MB107B), MBONs (MB011B, MB 298B) and pBD. Average bout length [H=19.23] and bout number [H=4.293] was analyzed by Kruskal Wallis ANOVA followed by Dunn’s post-hoc test. Bout length was elevated in MB010B, MB107B and MB011B. Bout number was consistent between genotypes and was no significantly different from control. (G) Activity or average beam counts/waking minute were consistent between flies expressing UAS-DopR2 RNAi (65997) in MB-KCs (MB010B, MB107B), MBONs (MB011B, MB 298B) and pBD. Activity was analyzed using one-way ANOVA [F (4,228) =3.818]. Post hoc Dunnett’s test reveals no significant differences in activity between tested genotypes and control.

As in the case of DopR1 receptor knockdown, daytime sleep (Figure 7B) was largely consistent between genotypes except MB298B where sleep was lower than control [F (4,228=65.45, p<0.0001)]. Increases in total sleep were largely contributed by nighttime sleep [F (4, 228) = 86.04, p<0.0001], Figure 7C and decreased sleep latency (Figure 7D) [F (4, 228) = 40.17, p<0.0001] indicating a consistent role for MB dopamine signaling via DopR1 and DopR2 in sleep initiation. In addition to decreased sleep latency, we also found that average length of sleep bout (Figure 7E) was higher in MB010B, MB107B and MB011B as compared to MB 298B and empty-pBD negative control [H=19.23]. Like DopR1, number of sleep bouts [H=4.293], Figure 7F and activity [F (4,228) =3.818], Figure 7G was consistent between genotypes.

Taken together, these results show that PAM dopamine signaling to specific MB compartments requires DopR1 and DopR2 receptor signaling specifically within the wake-regulating KCs or γ5 β’2 MBONs (MBON01,03 and 04) or both but not γ4 (MBON 05) compartment.

### 3.5 PAM γ5 signal via DopR1 and DopR2 to regulate total sleep and latency

While, UAS-RNAi transgene-induced gene silencing allows a spatial control, the efficacy of these transgenes and off-target effects are difficult to resolve in determining a clear role for DA receptors in MB mediated sleep regulation.

In order to directly address and test the coordinated role of PAM γ5 signaling through DopR1 and DopR2 receptor in wake regulation we specifically activated PAM γ5 (MB054B) neurons using a temperature sensitive cation channel dTrpA1 in DopR1 and DopR2 hypomorph backgrounds^62,64^. We measured total sleep in flies at 21°C (permissive temperature), the day before activation (baseline, Figure 8B) at which dTrpA1 channels are closed and the PAM γ5 (MB054B) neurons are inactive in w^1118^, DopR1, and DopR2 hypomorph background. We did not find any significant differences between the three tested genotypes during baseline [F (2,176) =2.815, p=0.08].

**Figure 8:**
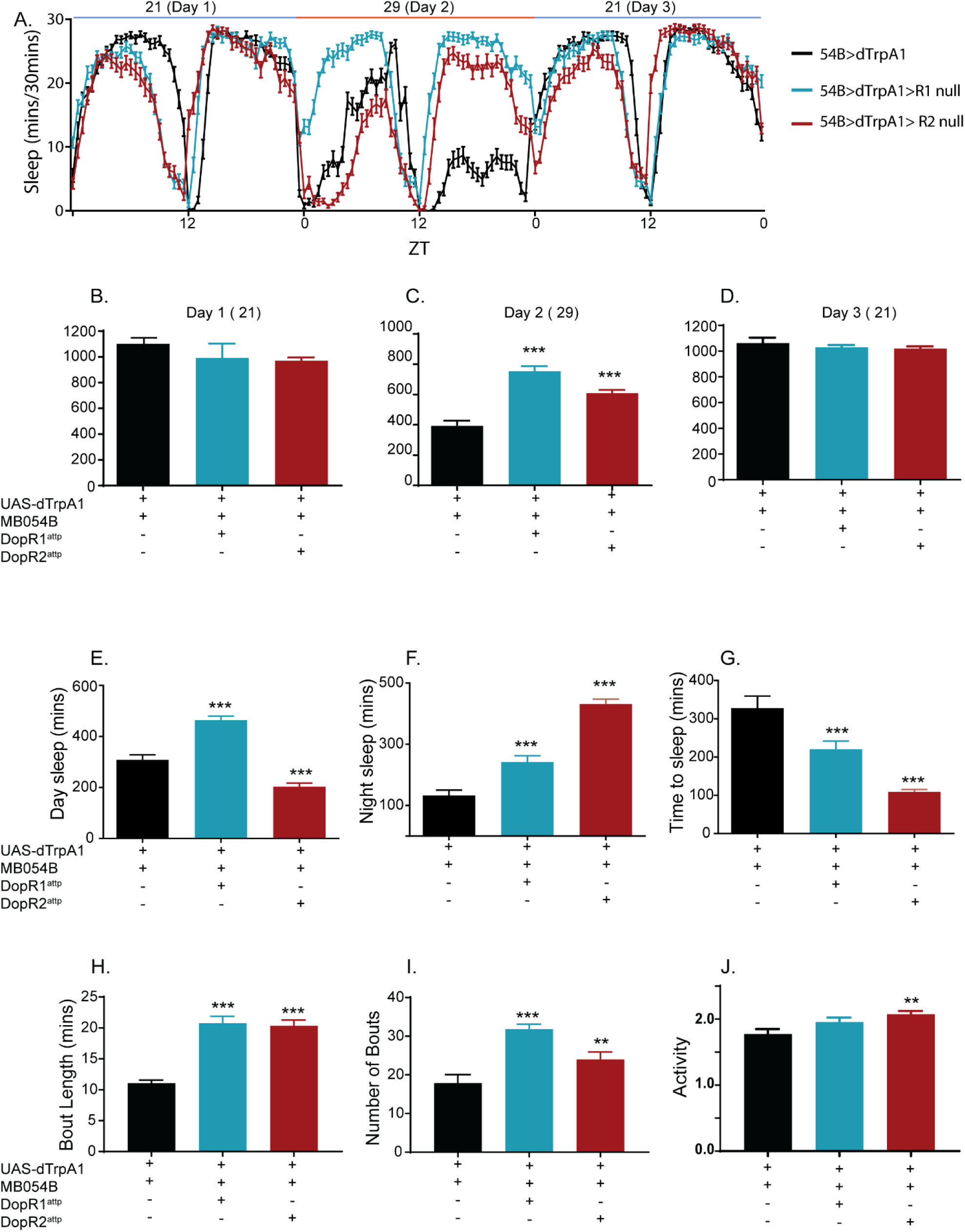
Wakefulness induced by PAM γ5 dopamine activation requires DopR1 and DopR2 function. (A) 3-day Sleep profile of flies expressing temperature sensitive cation channel dTrpA1 in PAM γ5 DANs (MB054B) in the wild type, DopR1 and DopR2 null backgrounds. Day 1 represents baseline at 21°C, Day 2 represent activation at 29°C and Day 3 represents recovery at 21°C. (B, C and D) Total sleep during Day 1 (baseline, 21°C), Day 2 (activation, 29°C) and Day 3 (recovery, 21°C), of flies expressing dTrpA1 in PAM γ5 in positive control (w^1118^) and receptor null/hypomorph background (DopR1^attp^ and DopR2^attp^). Total sleep was analyzed using one-way ANOVA for baseline [F (2,176) =2.815], activation [F (2,176) =19.6] and recovery [F (2,176) =1.832]. Post-hoc Dunnett’s test reveals no significant differences between PAM γ5 split-GAL4 in control and receptor null background in baseline and recovery condition. In activation condition (day 2) PAM γ5 activation in w^1118^ background induced wakefulness suppressed by receptor deficient background. For each of the experimental groups tested we had 59-61 flies which represented 2 independent experimental trials, sample included MB054B; dTrpA1/+ (59), MB054B; dTrpA1/DopR1^attp^ (59) and MB054B; dTrpA1/DopR2^attp^ (61) (E and F) Daytime and nighttime sleep during Day 2 (activation, 29°C) in flies expressing dTrpA1 in PAM γ5 in control and receptor null background (DopR1^attp^ and DopR2^attp^). One-way ANOVA for daytime [F (2,176) =76.20] and nighttime [F (2,176) =71.76] was followed by post-hoc Dunnett’s test. Daytime sleep was elevated in DopR1^attp^ and reduced in DopR2^attp^ background, while nighttime sleep was strongly elevated in both receptor null background. (G) Sleep latency or time to sleep (minutes) was found to be significantly reduced DopR1^attp^ and DopR2^attp^ background as compared to control background. One-way ANOVA for latency [F (2,176) = 24.41, p<0.0001] was followed by post-hoc Dunnett’s test. (H and I) Average bout length and bout number showed significant increases in receptor null background (DopR1^attp^ and DopR2^attp^) as compared to control. Average bout length [H=73.54] and bout number [H=52.39] was analyzed by Kruskal Wallis ANOVA followed by Dunn’s post-hoc test. (J) Activity or average beam counts/waking minute were consistent between flies expressing dTrpA1 in PAM γ5 in control and receptor null background (DopR1^attp^ and DopR2^attp^). One-way ANOVA for daytime [F (2,176) =3.868, p=0.02] and post-hoc Dunnett’s test revealed an increase in activity in DopR2^attp^ background

However, at 29°C when PAM neurons are activated in w^1118^ background we find significant decreases in sleep (~1000 minutes at day1/baseline to ~400 minutes on day 2/activation). Sleep suppression as a result of PAM activation affects both daytime and nighttime sleep, even though nighttime effects are stronger consistent with the role of PAM-DANs in nighttime sleep (Figure 8A). The sleep suppression caused by PAM γ5 activation was reversed or blocked in the DopR1 and DopR2 hypomorph background.

Wakefulness induced by PAM γ5 activation was blocked in the DopR1 and DopR2 background [F (2,176) =19.60, p<0.0001] and total sleep was significantly higher in flies where MB054B neurons are activated in DopR1 and DopR2 backgrounds as compared to w^1118^ (Figure 8C). These effects were reversible and total sleep is consistent between genotypes (Figure 8D), when temperature is switched back to 21°C (permissive temperature, Day 3), at which dTrpA1 is no longer open [F (2,176) =1.832, p=0.16].

The DopR1 and DopR2 hypomorphs had different effects on daytime and nighttime sleep induced by activation of PAM γ5 neurons. Both receptors strongly blocked nighttime sleep (Figure 8F) suppression caused by PAM activation [F (2, 176) = 71.76, p<0.0001]. Daytime sleep effects (Figure 8E) were differentially affected by the two receptor hypomorphs such that DopR1 hypomorph suppressed the waking effects of MB054B, while DopR2 did not [F (2, 176) = 76.2, p<0.0001]. The increased nighttime sleep was accompanied by a decrease in sleep latency (Figure 8G) consistent with RNAi mediated receptor manipulation of MBONs and KCs [F (2, 176) = 24.41, p<0.0001]. Waking activity at 29°C was consistent between tested genotypes [F (2, 176) = 3.868, p=0.023] with a small increase in DopR2 background (Figure 8J).

The wakefulness induced by activation of MB054B PAM DANs significantly reduces the bout length (H=73.54) and bout number (H=52.39) (Figure 8H and 8I). Both of these effects were blocked in the DopR1 and DopR2 hypomorph background suggesting that these receptors are required for regulation of sleep duration, latency and bout structure by PAM DANs.

## 4 Discussion

The mushroom body is a key structure involved in associative learning in insects^34,96-102^. The core-structure of the MB is identified by KC (Kenyon cell)-MBON (output neuron) synaptic compartments which are modulated by dopamine, octopamine, sNPF, GABA and SIFamide^37,56,103^. The mushroom body lobes are tiled by discrete anatomic compartments defined by the axons of a specific subset of DANs and the dendrites of one or two mushroom body output neurons (MBONs). This anatomical arrangement positions the DANs to strategically to convey positive and negative reinforced information by changing the synaptic weight of KC-MBONs in producing aversive and appetitive responses^37,56^.

While, the most in-depth analysis of these synapses and distinct DAN-KC-MBON connectivity and behavioral output comes from studies of olfactory conditioning, there is evidence that these synapses play a critical role in innate behaviors like feeding and sleep^3,41^.

A key defining feature of DAN modulation of MB involves lobe compartments where presynaptic arbors of the DANs and postsynaptic dendrites of the MBONs overlap defining the 15 compartmental units^37^. Our unbiased screen of PAM and PPL1 DANs projecting to all 15 compartments of the MB reveals that DAN inputs into both wake- and sleep-promoting KC-MBON microcircuits suppress sleep^3,33^. These wake-promoting MB-DANs were recently resolved by EM^104^ and project to γ5 (PAM01), γ4 (PAM 07,08), β’2amp (PAM02, PAM05 and PAM06), γ2α’1 (PPL1 103), α’3 (PPL1 104), α3 (PPL1 106) and α’2 α2 PPL1(PPL1 105). The KCs and MBONs downstream of PAM (01,02,05,06,07 and 08) can be neuroanatomically resolved and have been shown to be wake-promoting and alter their spontaneous neural activity in response to sleep-need (induced by mechanical sleep-deprivation). The ability to use EM connectivity in combination with cell-specific split-GAL4 tools provides opportunity to resolve the precise circuit mechanisms by which PAM neurons regulate wakefulness.

To understand the requirement of dopamine in MB-dependent sleep regulation we blocked release from sleep-regulating PAM and PPL1 DANs in the presence of a strong wake-inducing drug CBZ that blocks GABA transmission. GABA and Rdl based signaling within MB has been extensively studied in the context of associative learning and are thought to modulate KC and MBON activity via DANs^58,69,70^ but it is not clear if this regulation is generalized or specific to a few MB compartments. We find that CBZ does not directly change intracellular Ca++ levels in γ5 (PAM01), γ4 (PAM 07,08), β’2amp (PAM02, PAM05 and PAM06) neurons suggesting that either GABA mediated effects are downstream of PAM neurons or that the changes are small and undetectable with the CaLexA approach. While, it has been previously published that PAM neurons are broadly activated by caffeine using the CaLexA approach^78^, our studies show that caffeine effects are observable in γ5, γ4 and β’2 amp innervations. In addition to caffeine, the MB sleep-regulating compartment is also modulated by sleep need and thought to play a role in sleep homeostasis^33,105^. Our previous findings show that the γ5 β’2 MBONs and KCs show reduced activity post-sleep deprivation, indicating that their activity is negatively correlated to increased sleep drive during rebound period^33^. PAM01 (γ5) receives feedback activation from MBON γ5β’2a (MBON 01,03 and 04) mediated by NMDA receptors for the maintenance of short-term courtship memory^106^ and aversive memory extinction^49,79^. But unlike MBON γ5β’2a, we did not find evidence that PAM01 activity is altered by sleep-need.

A recent study using EM dataset of a Full Adult Female Fly Brain (FAFB) mapped the inputs and outputs of the PAMγ5 DANs and identified that this cell type is highly heterogenous and in addition to recurrent feedback from MBON01 γ5β’2a, it receives extensive input from other MBONs, sub-esophageal output neurons (SEZONs) and lateral horn output neurons^107^. The EM data also reveals that octopaminergic OA-VPM3 neurons synapse onto PAM01 (γ5), PAM01 (γ4) and weakly with PAM02 (β’2a) DANs. Whether, these inputs play a role in wakefulness is unknown but suggests that the PAMγ5 could serve as a key link between sensory inputs, wake-promoting octopamine signal and core sleep regulating circuitry within the MB. Each of these inputs could modulate PAM-DAN activity and dopamine release in regulating wakefulness via MB.

In addition to probing the release and activity of these PAM-DANs we also explored the dopamine receptors and their location within the MB in signaling wakefulness. To this end we expressed validated RNAi lines in subsets of KCs and MBONs and find that DopR1 and DopR2 are critical in mediating the wakefulness signal via KCs and γ5β’2 MBONs. The patterning of daytime and nighttime sleep is variable between tested genotypes but knocking down the receptor consistently increased total sleep by reducing latency and increasing bout length. Furthermore, specific manipulations of Dop1R receptors within MB did not directly alter waking activity as observed by manipulation of these receptors in CX^95^. The two Dop1R receptors regulate sleep in similar and potentially redundant way and are required for excitatory PAM-DAN signaling to MB γ5 and β’2 compartment. However, we did not find a role for these receptors in signaling wakefulness via the γ4 compartment.

In vitro characterization indicates Dop1R signal through distinct G-proteins, with DopR1 via Gαs to stimulate cAMP production^85,108^ and DopR2 coupling to Gαq via increased calcium^84,86^. These receptors have been studied extensively in the context of associative olfactory memory where DopR1 is necessary for memory formation, while DopR2 signaling is critical for erasing appetitive memories^93,109^. These receptors are thought to have differential sensitivity to dopamine^86^ and could be potentially recruited by varying DA release or DAN activity. Imaging activity in γ4 MBONs and KCs using recently developed GRAB_DA1m_ dopamine sensor^110^, cAMP^111^, and Ca^++112^ shows that these receptors are co-expressed and depending on the temporal relationship between an odor cue and dopaminergic reinforcement they can control the synapses bidirectionally^38^. This mechanism of producing different outcomes by differential temporal and spatial recruitment of receptors and downstream pathways in response to the same DANs are thought to underlie behavioral flexibility. In the context of sleep regulation, our work reveals that both DopR1 and DopR2 induce wakefulness in response to PAM γ5 and likely work synergistically in suppressing sleep.

Based on our results we posit a model (Figure 9) where PAM-DAN activity is not directly modulated by sleep-need or GABA signaling but activity of PAM-DANs is required for the wake effects of CBZ and reduced GABA transmission. Like PAM-DANs, previous studies show that MBON γ5β’2 are also required for wake effects of CBZ and their activity is directly altered by sleep and wake-need. Furthermore, MBONS and KCs that innervate the wake-active γ5 and β’2amp compartments require Dop1R’s to regulate sleep. From our RNAi and hypomorph experiments it appears that DopR1 knockdown has a stronger effect on sleep as compared to DopR2. But it’s not clear to what extend these receptors alter Ca++ and cAMP dynamics and how the downstream signaling affects behavioral output.

**Figure 9:**
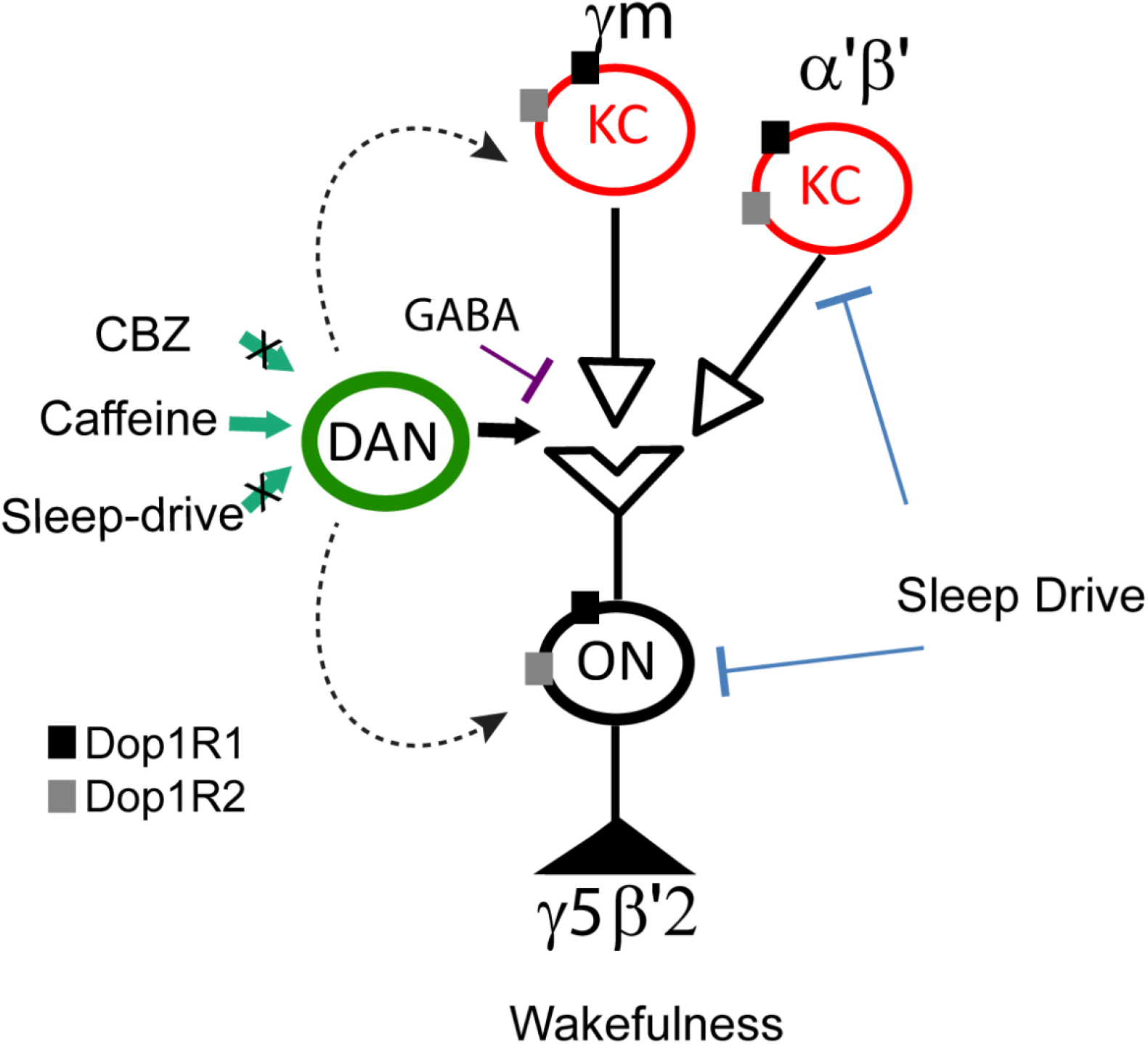
Schematic of PAM mediated sleep regulation within MBs. PAM-DANs projecting to KCs and MBONs of the γ5 and β’2 compartments of the MB signals wakefulness via DopR1 and DopR2 signaling. While the activity of the KCs and MBONS of these compartments is directly regulated by sleep-need, PAM-DAN activity is not. PAM-DAN mediated signaling is required for arousal/wakefulness induced by CBZ or altered GABA signaling.

The sleep-regulating PAM γ5 DANs and associated KCs and MBONs identified in our study are also involved in mediating satiety, novelty, punishment and reward associated experiences suggesting that the activity of these neurons is tuned to several wake and arousal associated behaviors. This is further supported by the EM connectome data showing that MB receives extensive gustatory, auditory and visual input in addition to olfactory input^104^.

Current models of sleep regulation rely on two main processes, the circadian clock and the sleep homeostat and don’t completely account for multiple external and internal factors that influence wakefulness^113^. The ability to sleep, however, is influenced by motivational or cognitive stimuli. Like MB-DANs, DA from ventral tegmental area (VTA) and projections to Nucleus Accumbens (NAc) in the midbrain plays a key role in processing reward and aversive signals and regulating cognition in rodents and primates^20^. Recent evidence from rodents shows that sleep and wakefulness are regulated by VTA DANs of the mesolimbic pathway^21^. Indeed, like PAM-DANs, VTA-dopaminergic neuron activation is sufficient to drive wakefulness and necessary for the maintenance of wakefulness and inhibition of VTA DANs promoted sleep specifically in the presence of wake-inducing stimuli and these effects are mediated via NAc^5,21,114,115^. The dopamine signaling from VTA to NAc co-regulates sleep with other behaviors in a context dependent manner like the MB, but how this co-regulation is achieved requires further study.

We therefore envision that sleep, wakefulness and arousal within MB are not located in distinct circuits, but rather mediated by distinct processes within a common circuit. Hence, encoding distinct activity phases within a common circuit may be an efficient mechanism for integrating long and short-term suppression of sleep to coordinate motivated processes with wakefulness.

## Acknowledgments

We thank Dr. Gerald Rubin, Dr. Yoshinori Aso and Dr. Jing Wang and for generously providing fly stocks. Several transgenic lines were obtained from Bloomington Drosophila Stock Center. We also thank Dr. Callen Hyland, Lilian Mworia, Catherine Shorb, Zani Moore, Pomai Murakami and Austin Pavin for help with the experiments and aspects of fly rearing, behavioral set-ups and confocal imaging. The project was supported by NIH grant 1R15GM125073-01 awarded to Dr. Divya Sitaraman and student research awards to Steven Buchert, Margaret Driscoll, Morgan McLaughlin and Amanda Nguyen.

## Data Availability Statement

The data that support the findings of this study are available on request from the corresponding author.

---

**Figure S1:**
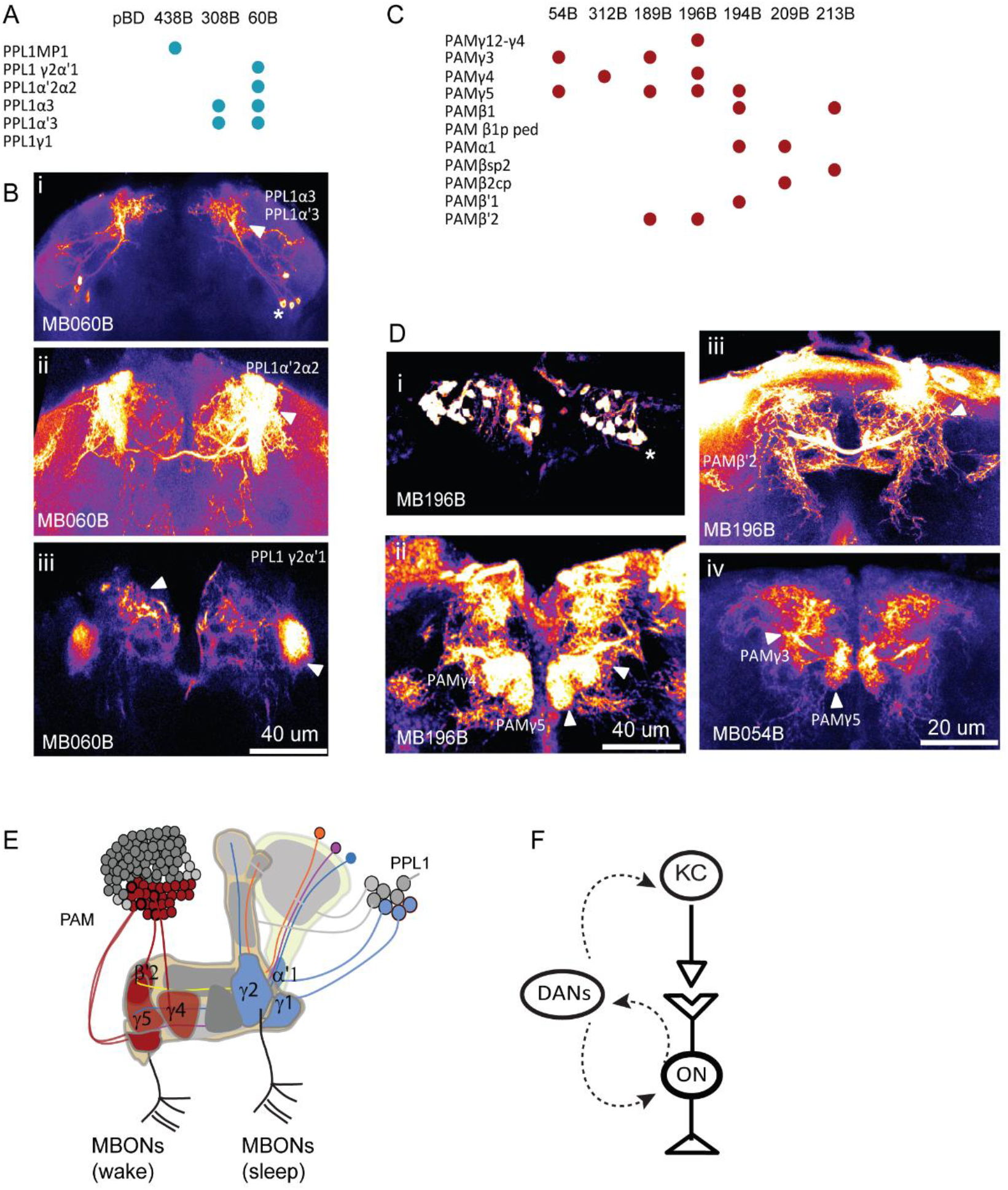
Lobe specific innervations of PAM and PPL1 DAN subsets. The expression patterns with lobe specific innervation of these specific and broad sleep-regulating dopamine drivers was confirmed by expression of UAS-mCD8-GFP. (A and C) Lobe specific expression of MB-DANs split-GAL4 lines MB060B (broad PPL1), MB196B (broad PAM) and MB054B (PAM subsets). (B and D) Maximum intensity projections of confocal stacks (10-15 slices) representing the MB regions in fire LUT applied to emphasize PPL1 (B i-iii) and PAM (D i-iv) neurons. Cell bodies (asterisks) and lobe specific innervations (arrowhead) have been indicated to identify MB compartment innervated by each of the split-GAL4 used. Scale bars indicate 20um or 40um. (E) Schematic representation of MB showing lobe specific axonal projections of PAM (red) and PPL1 (blue) DANs. Subset of PAM innervate the MB compartments (red) involved in regulating wakefulness mediated by KCs and MBONs. (F) Schematic representation of synaptic interactions between PAM/PPL1 DANs, KCs and MBONs.

**Figure S2:**
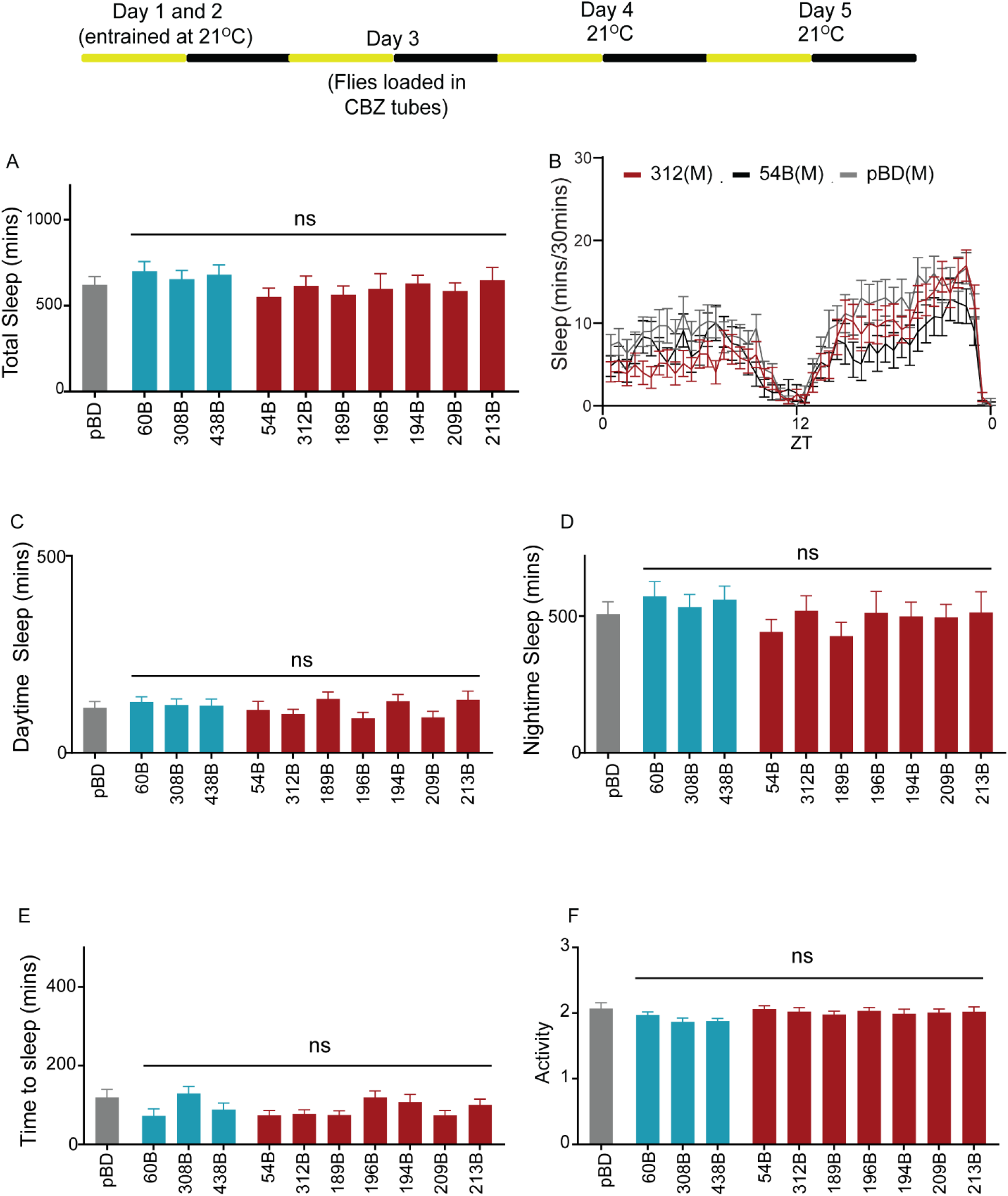
Inhibition of PAM and PPL1 DANs but not genotypic differences promote sleep in CBZ treated flies. Schematic of experimental protocol showing temperature and drug conditions over a 5-day period. Mated female flies were loaded on 0.1mg/ml CBZ containing food (Day 3) in incubator maintained at 21°C. Day 4 the temperature was maintained at 21°C and sleep was measured using the Drosophila Activity Monitoring system. (A) Total sleep in MB-DANs (PPL1-blue: MB060B, MB308B, MB438B and PAM-red: MB054B, MB312B, MB194B, MB196B, MB209B, MB213B, MB189B) expressing temperature sensitive dominant negative dynamin mutation, Shi^ts1^ in the presence of CBZ (Day 4). (B) Representative sleep profile of flies expressing Shi^ts1^ specific PAM (MB054B-black and MB312B-red) neurons with empty/enhancerless-GAL4 control (pBD, grey). (C) Daytime sleep during the first 12-hour period of specific PPL1 (blue), PAM (red) neurons as compared to empty-GAL4 control (pBD, grey). (D) Nighttime sleep during of specific PPL1 (blue), PAM (red) neurons as compared to empty-GAL4 control (pBD, grey). Statistical significance of total sleep, daytime and nighttime sleep was determined by performing one-way ANOVA followed by Dunnett post-hoc analysis. For total sleep [F (10, 469) =0.7056, p=0.7195], daytime sleep [F (10, 469) =0.8935, p=0.5391] and nighttime sleep [F (10, 469) =0.7018, p=0.7230]. For each of the experimental groups we had 37-52 flies which represented 2 independent experimental trials. Specifically, we tested pBD (n=45), 54B (n=46), 60B (n=37), 189B (n=45), 194B (n=45), 209B (n=45), 213B (n=38), 308B (n=51), 312B (n=40), 438B (n=48) and 196B (n=40). (E) Sleep latency or time to sleep was calculated as the time gap between lights off and first sleep bout and analyzed using one-way ANOVA [F (10, 469) =1.240, p=0.263] followed by post-hoc Dunnett test. (F) Activity or average beam counts/waking minute were consistent between PAM, PPL1 and controls [F (10,469) =1.757, p=0.06]

**Figure S3:**
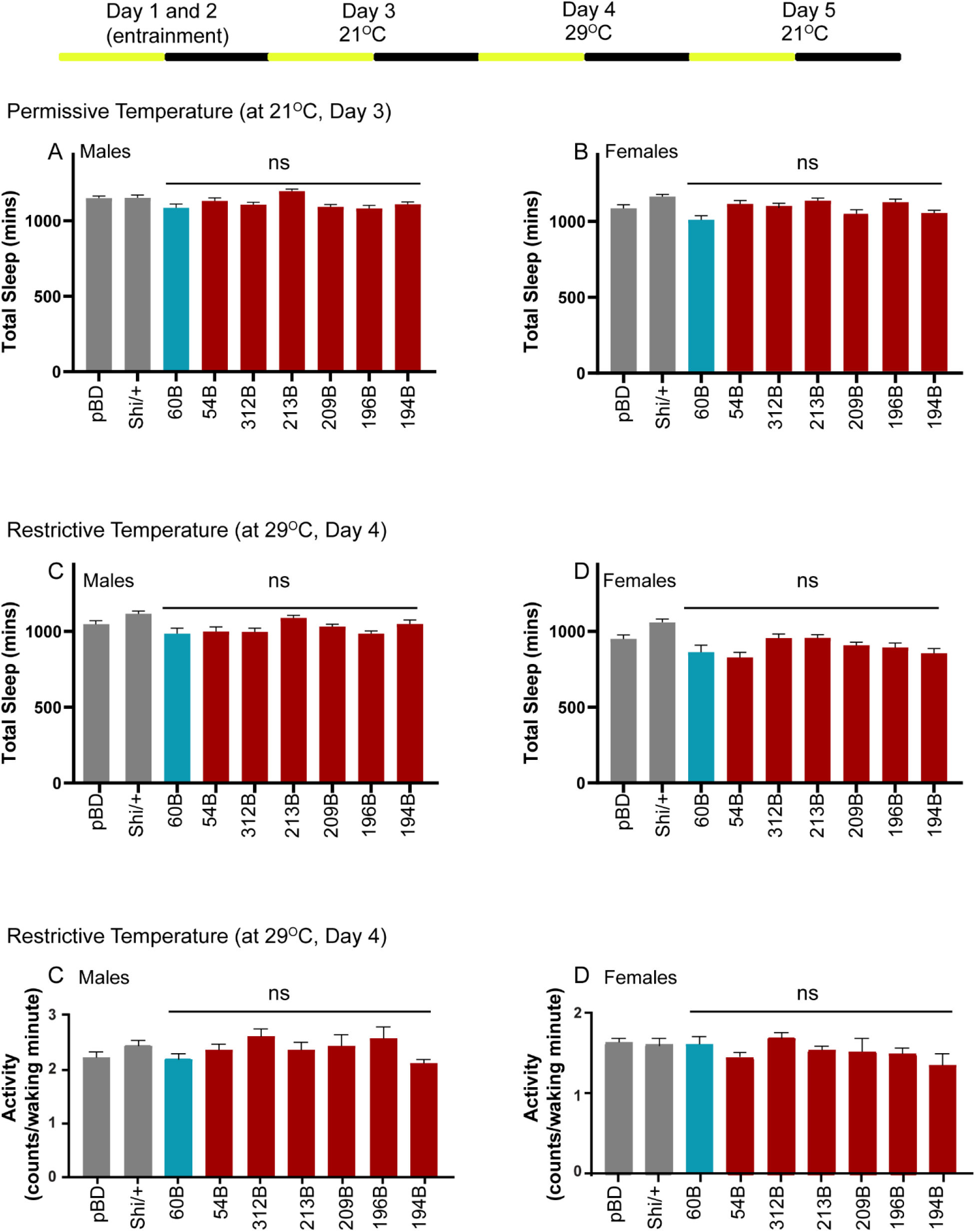
PAM and PPL1 inhibition does not increase sleep over baseline in the absence of CBZ in males and females. Schematic of experimental protocol showing temperature and drug conditions over a 5-day period. Male and female flies were loaded in 5% sucrose and 2% agarose containing food and sleep was measured post-entrainment on Day 3 (21°C-permissive temperature) and Day 4 (29°C-permissive temperature). In addition to pBD we also tested Shi/+ flies as an additional negative control. (A and B) Total sleep in males and females with MB-DANs (PPL1-blue: MB060B and PAM-red: MB054B, MB312B, MB196B, MB209B, MB213B and MB194B) expressing Shi^ts1^ in the absence of CBZ (Day 3, 21°C). Total sleep was analyzed using one-way ANOVA in males [F (8,233) =4.35] and females [F (8,241) =3.79]. Post-hoc Dunnett test in males and females did not detect significant differences between tested genotypes and controls. For each of the experimental groups tested we had 20-32 flies which represented 2 independent experimental trials. For males the sample included pBD (n=29), 54B (n=32), 60B (n=28), 189B (n=22), 194B (n=20), 209B (n=28), 213B (n=24), 312B (n=32) and 196B (n=27). For female flies the sample included pBD (n=28), 54B (n=30), 60B (n=32), 189B (n=26), 194B (n=22), 209B (n=31), 213B (n=25), 312B (n=29) and 196B (n=31). (C and D) Total sleep in males and females with MB-DANs (PPL1-blue: MB060B and PAM-red: MB054B, MB312B, MB196B, MB209B, MB213B and MB194B) expressing Shi^ts1^ in the absence of CBZ (Day 4, 29°C). Total sleep was analyzed using one-way ANOVA in males [F (8,233) =4.416] and females [F (8,241) =5.357]. Post-hoc Dunnett test in males and females did not detect significant differences between tested genotypes and controls. (E and F) Activity in males and females with MB-DANs (PPL1-blue: MB060B and PAM-red: MB054B, MB312B, MB196B, MB209B, MB213B and MB194B) expressing Shi^ts1^ in the absence of CBZ (Day 4, 29°C). Activity was analyzed using one-way ANOVA in males [F (8,233) =1.457, p=0.1742] and females [F (8,241) =1.291, p=0.249]. Post-hoc Dunnett test in males and females did not detect significant differences between tested genotypes and controls.

**Figure S4:**
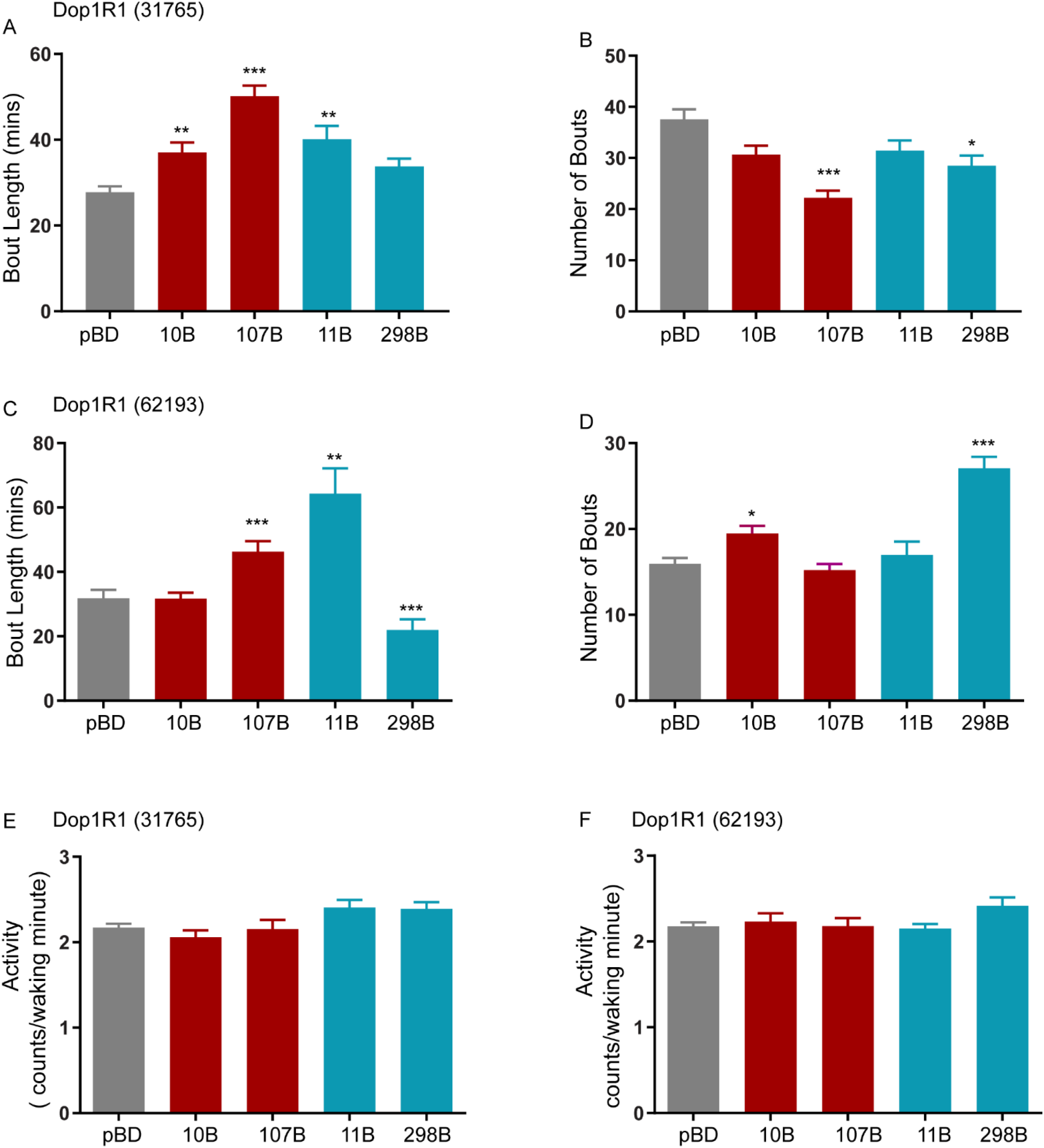
Knockdown of DopR1 MB-KCs and MBONs increases sleep by changing bout length but does not affect waking activity. (A and B) Average bout length and bout number in flies expressing UAS-DopR1 RNAi (31765) in MB-KCs (MB010B, MB107B), MBONs (MB011B, MB 298B) and pBD. Average bout length [H=50.28] and bout number [H=32.97] was analyzed by Kruskal Wallis ANOVA followed by Dunn’s post-hoc test. Bout length was elevated in M010B, MB107B and MB011B. Bout number was significantly reduced in MB107B and MB298B. For each of the experimental groups tested we had 37-48 flies which represented 2 independent experimental trials, sample included 31765/pBD (48), 31765/MB010B (40), 31765/MB107B (38),31765/MB011B (38),31765/MB298B (37). (C and D) Average bout length and bout number in flies expressing UAS-DopR1 RNAi (62193) in MB-KCs (MB010B, MB107B), MBONs (MB011B, MB 298B) and pBD. Average bout length [H=48.56] and bout number [H=49.70] was analyzed by Kruskal Wallis ANOVA followed by Dunn’s post-hoc test. Bout length was elevated in MB107B and MB011B and reduced for MB298B. Bout number was significantly increased in MB010B and MB298B. For each of the experimental groups tested we had 39-47 flies which represented 2 independent experimental trials, sample included 62193/pBD (47), 62193/MB010B (39), 62193/MB107B (44), 62193/MB011B (43), 62193/MB298B (47). (E and F) Activity or average beam counts/waking minute were consistent between flies expressing UAS-DopR1 RNAi (62193 and 31765) in MB-KCs (MB010B, MB107B), MBONs (MB011B, MB 298B) and pBD. Activity was analyzed using one-way ANOVA for 31765 [F (4,196=3.424)] and for 62193 [F (4,220=0.2261)]. Post hoc Dunnett’s test reveals no significant differences in activity between tested genotypes and control.

